# Melanocortin-4 receptor antagonist TCMCB07 alleviates chemotherapy-induced anorexia and weight loss

**DOI:** 10.1101/2024.09.14.613069

**Authors:** Xinxia Zhu, Russell Potterfield, Kenneth A. Gruber, Emma Zhang, Samuel Newton, Mason A. Norgard, Peter R. Levasseur, Peng Bai, Xu Chen, Qingyang Gu, Aaron J. Grossberg, Daniel L. Marks

## Abstract

Cancer patients undergoing chemotherapy often experience anorexia and weight loss that significantly deteriorates overall health, reduces treatment tolerance and quality of life, and worsens oncologic outcomes. There are currently few effective therapeutic options to mitigate these side effects. The central melanocortin system, which plays a pivotal role in regulating appetite and energy homeostasis, presents a logical target for treating anorexia and weight loss. In this preclinical study, we evaluated the efficacy of TCMCB07, a synthetic antagonist of the melanocortin-4 receptor, in mitigating anorexia and weight loss in several rat models of chemotherapy: cisplatin, 5-fluorouracil, cyclophosphamide, vincristine, doxorubicin, and a combination of irinotecan and 5-fluorouracil. Our results indicate that peripheral administration of TCMCB07 improved appetite, stabilized body weight, preserved fat and heart mass, and slightly protected lean mass after multiple cycles of chemotherapy. Furthermore, combining TCMCB07 with a growth differentiation factor 15 antibody enhanced treatment effectiveness. Similar effects from TCMCB07 treatment were observed in a rat tumor model following combination chemotherapy. No significant adverse effects nor increased chemotherapy-related toxicities were observed with TCMCB07 treatment. These findings suggest that peripheral administration of TCMCB07 holds promise as a therapeutic approach for alleviating chemotherapy-induced anorexia and weight loss, potentially benefiting numerous patients undergoing chemotherapy.

**Graphical abstract:** 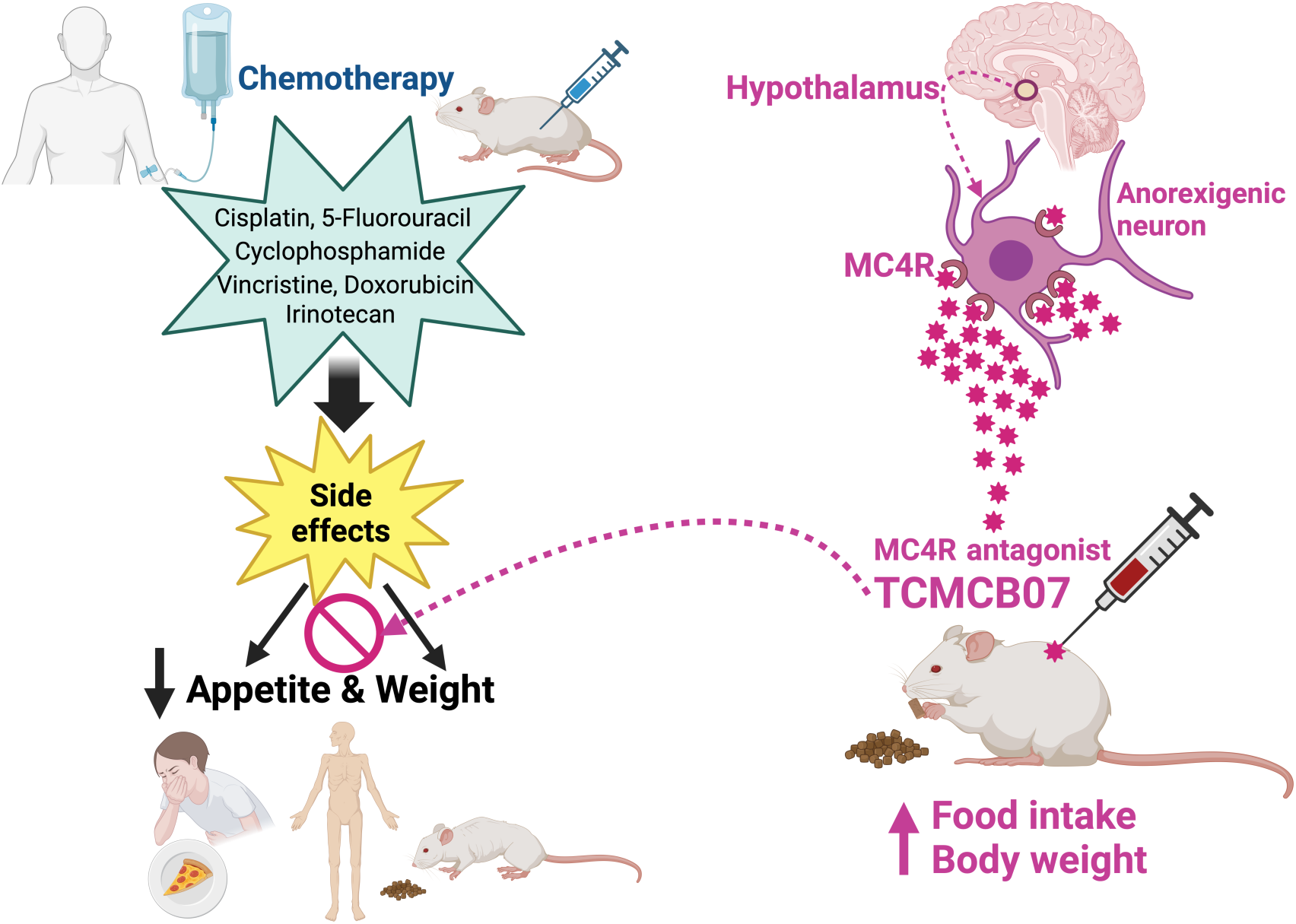

## Introduction

Chemotherapy remains a cornerstone of both curative and palliative cancer therapy. It is the primary approach for treating advanced malignancies in situations where surgical resection or radiation therapy is not viable (1, 2). Chemotherapy delivery has been improved by refining dosing strategies, incorporating neoadjuvant or adjuvant administration, and integration with more advanced supportive care (1). However, because cancer cells are so similar to healthy cells, delivering adequate cytotoxic doses often results in pronounced adverse effects like nausea, vomiting, anorexia, and weight loss. These side effects can profoundly impact patients’ quality of life and limit treatment adherence (3–5). Therefore, effective management of chemotherapy-induced side effects is imperative for both patients and clinicians.

In recent decades, there have been substantial improvements in management of chemotherapy-induced nausea and vomiting through the utilization of standard-of-care agents, such as 5-hydroxytryptamine 3 receptor (5-HT3) antagonists, neurokinin-1 receptor (NK-1) antagonists, dexamethasone, and olanzapine (6–8). However, a significant number of individuals undergoing chemotherapy experience decreased appetite and weight loss, which not only deteriorates their overall physical condition, but also reduces treatment tolerance and exacerbates the underlying disease. Recent research revealed circulating levels of GDF15 are elevated by chemotherapies (9). GDF15 triggers nausea, vomiting, anorexia, and weight loss via activating glial cell-derived neurotrophic factor receptor alpha-like (GFRAL) in the area postrema and nucleus of the solitary tract in the brainstem and other GDF15 receptors or pathways (10–14). A recent study reported that treatment with a GDF15 neutralizing antibody mitigated cisplatin-induced vomiting, anorexia, and weight loss, indicating the potential of this therapeutic approach for alleviating side effects induced by platinum-based chemotherapy (15).

The central melanocortin system plays a pivotal role in the regulation of appetite, body weight, and energy homeostasis. This system encompasses first-order orexigenic agouti-related peptide (AgRP) neurons and anorexigenic proopiomelanocortin (POMC) neurons in the arcuate nucleus of hypothalamus, which sense hormonal (ghrelin, leptin, and insulin) and neuronal (excitatory and inhibitory inputs) signals of energy balance and activate downstream neurons (16–18). These neurons synapse on second-order neurons in the paraventricular nucleus that express the melanocortin 3 receptor (MC3R) and MC4R, neuropeptide Y receptor 1, and GABA_A_ receptors, and are therefore capable of integrating inputs from AgRP and POMC neurons (19). These neurons transduce both anorexigenic agonists (e.g., alpha-melanocyte-stimulating hormone [α-MSH]) and orexigenic antagonists/inverse agonists (e.g., AgRP) of MC3R and MC4R. While MC3R neurons likely contribute to behavioral adaptation to fasting and nutrient partitioning, MC4R neurons are involved in feeding behavior, adaptive thermogenesis, and glucose homeostasis (20, 21). Therefore, this system represents a rational therapeutic target for treating anorexia, cachexia, obesity, and diabetes (19, 22–24).

Over the last decade, the development of melanocortin receptor-based therapeutics brought excitement and promise in treating obesity and anorexia-cachexia, with several melanocortin agonist drugs approved by the FDA for certain forms of obesity and other neuroendocrine disorders (19). However, despite investigations into melanocortin antagonists for therapeutic interventions to treat wasting syndromes and anorexia, no drugs in this class are yet approved for clinical use (25–30), highlighting the need to develop novel drugs with maximum safety, high efficacy, and peripheral therapeutic feasibility (e.g., effectively crossing the blood-brain barrier [BBB] to act on the central melanocortin system). Our previous work demonstrated the efficacy of TCMCB07, a synthetic MC4R antagonist, in ameliorating cachexia associated with cancer, chronic kidney disease, and other advanced illnesses (31–35). TCMCB07 recently completed a first-in-human phase 1 clinical trial, with preliminary findings supporting both safety and efficacy (36).

In this preclinical study, we aimed to investigate the potential of TCMCB07 in alleviating chemotherapy-induced anorexia and weight loss. We replicated chemotherapy-associated anorexia and weight loss in rats by administering six commonly used cytotoxic agents, either individually or in combination: cisplatin, 5-fluorouracil (5-FU), cyclophosphamide, vincristine, doxorubicin, and irinotecan. We also utilized the rat Ward colorectal carcinoma model in combination with chemotherapy to model chemotherapy treatment of a tumor *in situ* (37–41). A comprehensive evaluation of TCMCB07’s efficacy in stimulating appetite and maintaining body mass was performed across these rat models. Additionally, we explored the possibility of enhancing the therapeutic effectiveness by combining TCMCB07 with an anti-GDF15 antibody to combat chemotherapy-induced anorexia and weight loss.

## Results

### Dosing regimen selection for TCMCB07 administration and chemotherapy

We selected the dose of TCMCB07 based on our prior work demonstrating that 3 mg/kg/day effectively ameliorates cachexia associated with cancer, renal failure, or other advanced conditions (33–35). To assess the broad effectiveness of TCMCB07 treatment in chemotherapy-induced anorexia and weight loss, we generated six rat models of chemotherapy, representing commonly used classes of cancer chemotherapeutics: 1) cisplatin (platinum compound), 2) 5-FU (antimetabolite), 3) cyclophosphamide (alkylating agent), 4) vincristine (vinca alkaloid), 5) doxorubicin (anthracycline), and 6) combination of irinotecan (DNA topoisomerase I inhibitor) and 5-FU. A literature review of chemotherapy-induced anorexia and weight loss revealed a wide range of chemothersapy doses (42–48). Therefore, we designed and conducted a series of dose-response experiments in rats to determine doses that consistently mimicked the commonly observed clinical sickness responses, without undue toxicity. Initially, two doses of cisplatin and 5-FU were tested: cisplatin at 2.5 and 5 mg/kg, and 5-FU at 62.5 and 125 mg/kg (Supplemental Figure 1. A-C). For cyclophosphamide and vincristine, four doses of each agent were examined: cyclophosphamide at 50, 70, 90, and 110 mg/kg (Supplemental Figure 1, D and E), and vincristine at 0.18, 0.25, 0.30, and 0.40 mg/kg (Supplemental Figure 1, F and G). For doxorubicin, a dosage of 2 mg/kg was chosen based on previous reports (49, 50). For the combination of irinotecan and 5-FU, a dosage of 50 mg/kg for each agent was selected following previous studies (37–41). All chemotherapy agents were administered via IP injections once per week for three cycles in total. Control animals received an equivalent volume of saline IP injections. Through these dose-response experiments, we observed not only dose-dependent general behavioral responses to chemotherapy, such as reduced activity indicating fatigue, but also a gradual decline in food intake and body weight over multiple cycles of chemotherapy, consistent with common clinical side effects induced by these drugs. In addition, we noted severe morbidity and even mortality among animals receiving high doses of each chemotherapy agent. Based on these results and considering the maximum tolerance of animals to three consecutive cycles of chemotherapy plus daily TCMCB07 administration, we ultimately selected the optimal dose for each chemotherapy agent to induce a 10 to 30% weight loss compared to rats not receiving chemotherapy (15). Rats were treated for a total of three consecutive cycles (administered once per week, via IP injection) at the following doses (in mg/kg): cisplatin 2.5, 3.0, or 5.0 (with the dose of 5.0 mg/kg administered for the first cycle only), 5-FU 70, cyclophosphamide 65, vincristine 0.27, doxorubicin 2.0, and irinotecan 50 and 5-FU 50 for two cycles of combination chemotherapy.

### TCMCB07 treatment restores appetite during multiple cycles of chemotherapy

We conducted independent studies to evaluate TCMCB07’s efficacy across the five rat models of single-agent chemotherapy (Figure 1A): 1) cisplatin, 2) 5-FU, 3) cyclophosphamide (CP), 4) vincristine, and 5) doxorubicin. In the cisplatin/TCMCB07 study, during three cycles of chemotherapy, we observed a significant attenuation of chemotherapy-induced anorexia and a faster rebound in daily food intake among rats treated with TCMCB07 compared to those receiving saline treatment (Figure 2A). Cumulative food intake over the 21-day study period was identical between cisplatin/TCMCB07 and saline/saline groups, indicating that TCMCB07 treatment completely reversed the anorexia induced by multiple cycles of cisplatin chemotherapy (Figure 2B). Similarly, in the 5-FU/TCMCB07 study, TCMCB07 treatment led to increased cumulative food intake compared to both saline/saline and 5-FU/saline groups by day 21 (Figure 2, C and D). In the cyclophosphamide/TCMCB07 and vincristine/TCMCB07 studies, although TCMCB07 treatment did not elevate food intake during the initial four days following the first chemotherapy treatment, it increased daily food intake for the remainder of the study period. (Figure 2, E and G). TCMCB07 treatment reversed any chemotherapy-induced reduction in cumulative food intake in both the CP and vincristine-treated groups (Figure 2, F and H). Following each cycle of cisplatin or CP chemotherapy, weekly food intake was significantly greater in rats receiving TCMCB07 compared to those receiving saline (Figure 2, I and K). During the second and third cycle of 5-FU or vincristine chemotherapy, weekly food intake was higher in rats receiving TCMCB07 compared to those receiving saline (Figure 2, J and L). Notably, no significant reduction in weekly food intake was observed in any of the chemotherapy/TCMCB07 groups compared to saline/saline groups (Figure 2, I-L). Food intake was not measured in the doxorubicin/TCMCB07 study. Taken together, peripheral administration of TCMCB07 completely reversed anorexia during three cycles of cisplatin, 5-FU, cyclophosphamide, or vincristine chemotherapy, suggesting TCMCBC’s efficacy in restoring appetite suppressed by these commonly used chemotherapy agents.

**Figure 1:**
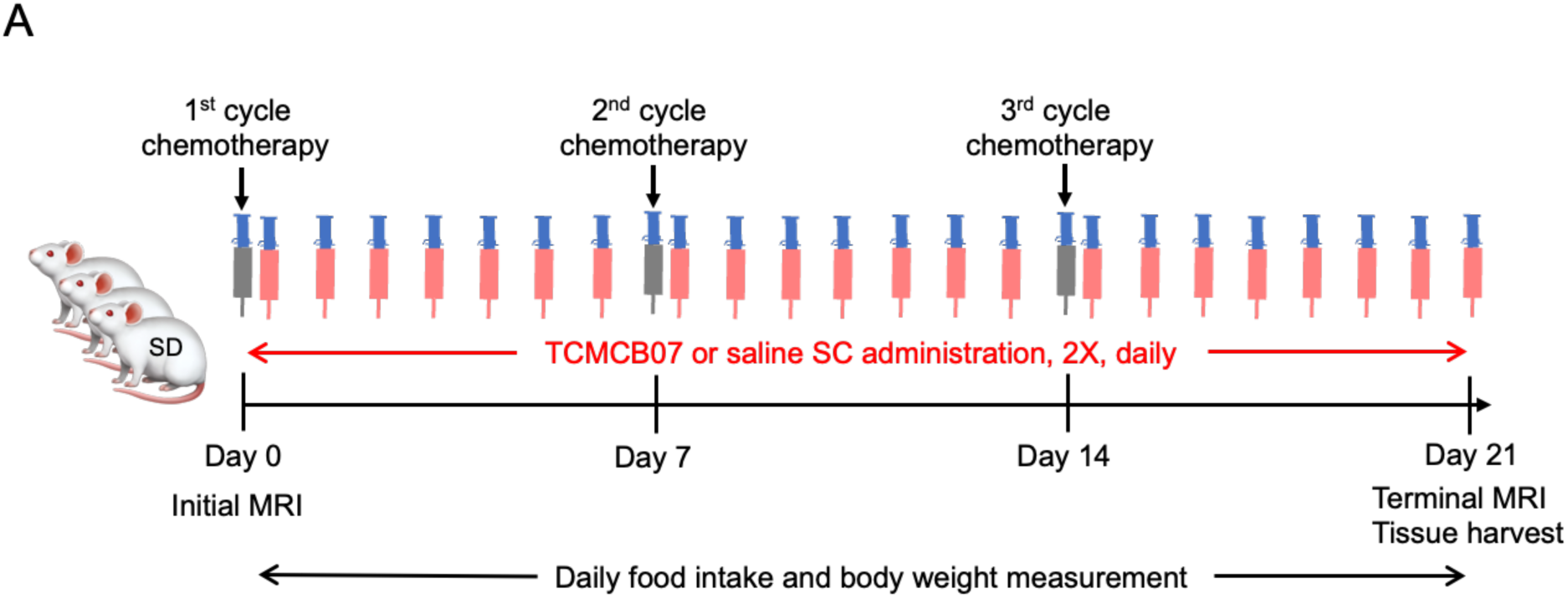
Schematic of TCMCB07/chemotherapy study design. (**A**) Schematic of TCMCB07/chemotherapy study design. Sprague-Dawley (SD) male rats were treated with three cycles of chemotherapy via IP injection within three weeks at doses: cisplatin 2.5 mg/kg, 5-Fluorouracil 70 mg/kg, cyclophosphamide 65 mg/kg, vincristine 0.27 mg/kg, and doxorubicin 2 mg/kg. Control animals received an equivalent volume of saline IP injections. All rats received SC injections twice (2x) daily with either saline or TCMCB07 (3 mg/kg/day) from day 0 to 21. Initial and terminal body composition was measured using MRI prior to and post treatments. Food intake and body weight were monitored daily throughout entire experimental period (day 0-21). At the end of the experiment, tissues were harvested following euthanasia.

**Figure 2:**
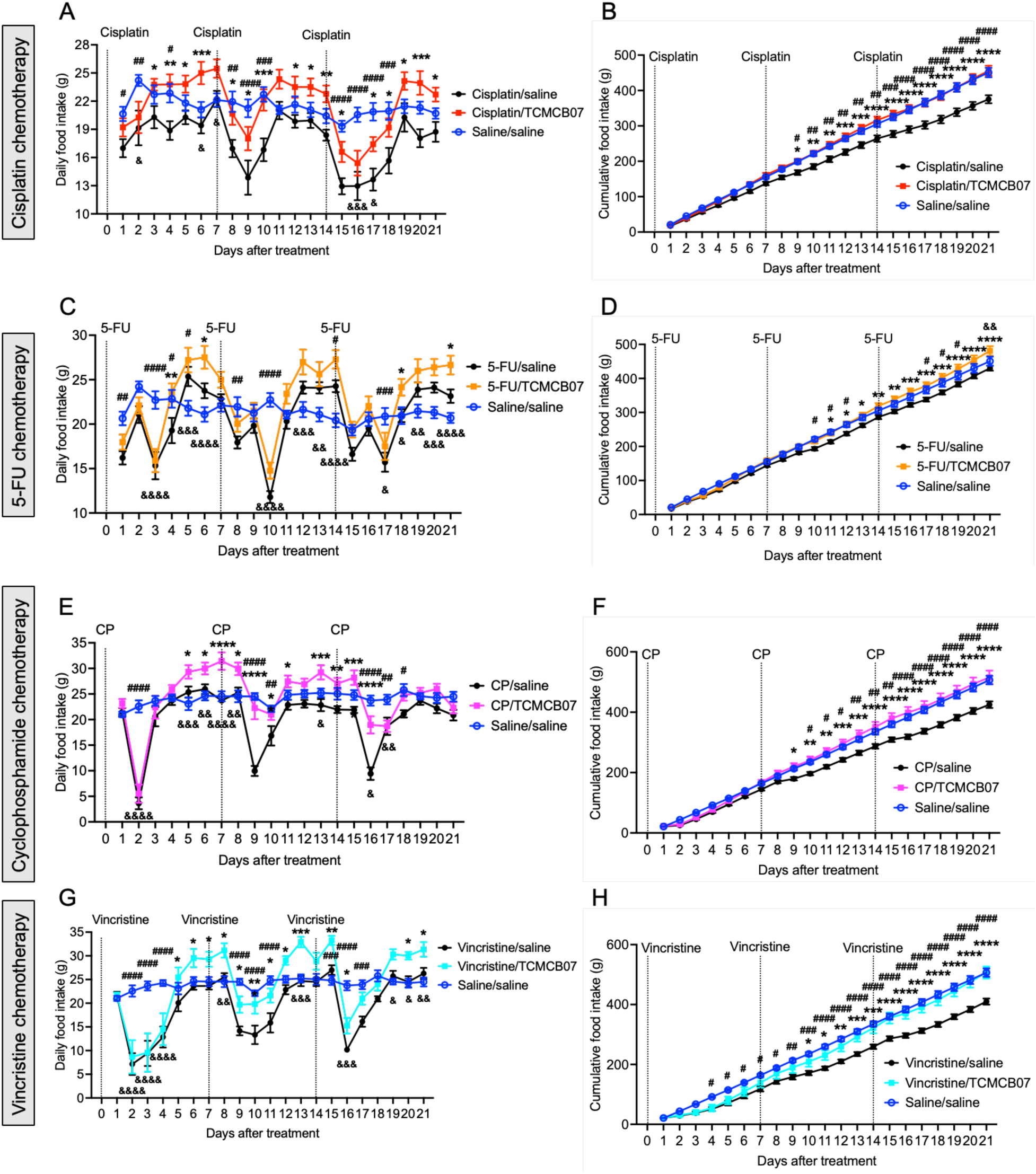
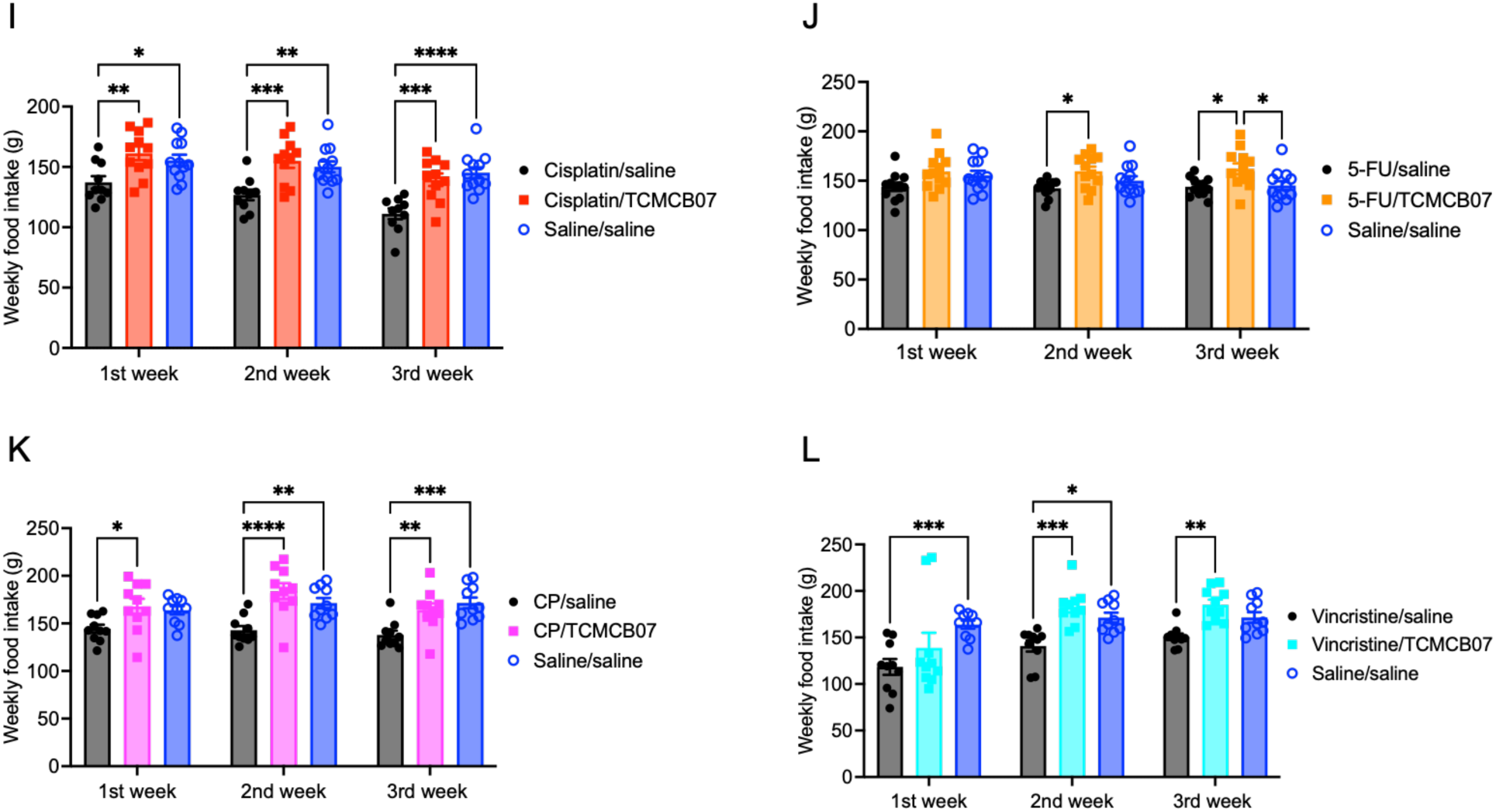
**TCMCB07 treatment restores appetite during multiple cycles of chemotherapy. (A-H**) Daily and cumulative food intake after chemotherapy and TCMCB07 treatment throughout entire experimental period (day 0-21). (**A**) and (**B**) Cisplatin chemotherapy. (**C**) and (**D**) 5-Fluorouracil chemotherapy. **(E**) and (**F**) Cyclophosphamide chemotherapy. **(G**) and (**H**) Vincristine chemotherapy. (**I-L**) Weekly food intake after cisplatin (**I**), 5-Fluorouracil (**J**), cyclophosphamide (**K**), or vincristine (**L**) chemotherapy and TCMCB07 treatment. All data in (**A**-**H**) are expressed as mean ± SEM for each group, and all data in (**I**-**L**) are expressed with each dot representing one sample. *n* = 10-12. (**A**-**H**), *: Chemotherapy/saline vs Chemotherapy/TCMCB07, #: Chemotherapy/saline vs Saline/saline, &: Chemotherapy/TCMCB07 vs Saline/saline. *, #, &, *P* < 0.05; **, ##, &&, *P* < 0.01; ***, ###, &&&, *P* < 0.001; ****, ####, &&&&, *P* < 0.0001. All data in (**A-H**) are analyzed by Two-way ANOVA, and all data in (**I-L**) are analyzed by One-way ANOVA.

### TCMCB07 treatment preserves body mass throughout multiple cycles of chemotherapy

In the same experiments outlined above, we monitored the daily body weight in the same groups of rats. Following each cycle of cisplatin chemotherapy, rats experienced weight loss compared to those not receiving chemotherapy, with the loss becoming more pronounced after second and third cycles of chemotherapy (Figure 3A, Supplemental Figure 2A). TCMCB07 treatment completely reversed this weight loss compared to saline treatment, as there was no difference in body weight between the cisplatin/TCMCB07 and saline/saline groups at the end of each cycle of cisplatin chemotherapy (Figure 3B). Similarly, while rats receiving 5-FU experienced weight loss compared to those receiving saline (Figure 3C, Supplemental Figure 2B), TCMCB07 treatment fully mitigated this weight loss during the three cycles of chemotherapy (Figure 3D). Although TCMCB07 treatment did not completely reverse the body weight loss induced by cyclophosphamide or vincristine chemotherapy, it significantly attenuated the reduction in body weight, particularly during the second and third cycles of chemotherapy (Figure 3, E-H, Supplemental Figure 2, C and D). With doxorubicin chemotherapy, TCMCB07 treatment alleviated body weight loss compared to saline treatment during the three cycles of chemotherapy (Figure 3, I and J, Supplemental Figure 2E). Collectively, TCMCB07 treatment attenuated body weight loss induced by three cycles of cisplatin, 5-FU, cyclophosphamide, vincristine, or doxorubicin chemotherapy, suggesting the efficacy of TCMCB07 in preserving body mass during multiple cycles of chemotherapy.

**Figure 3:**
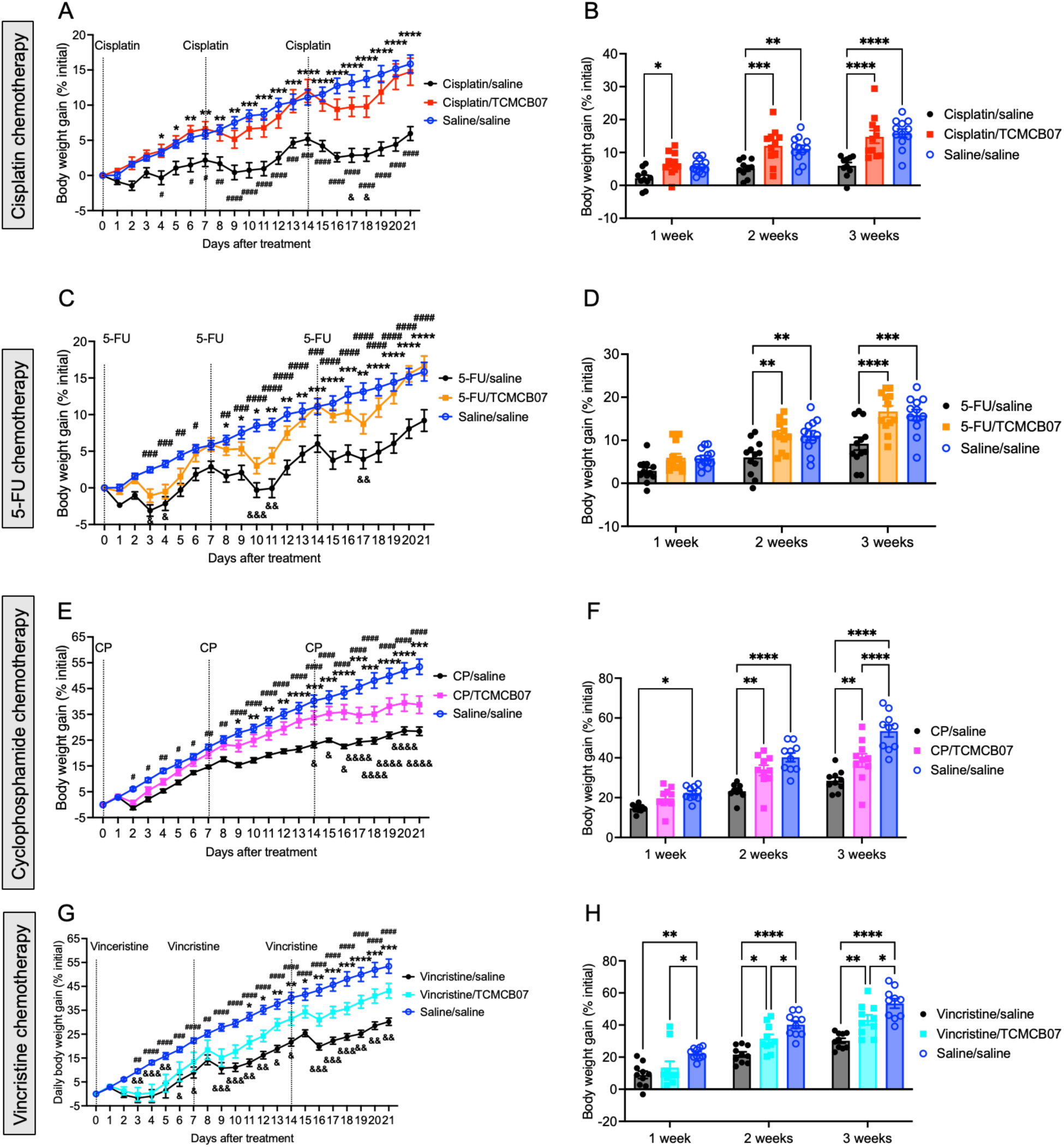
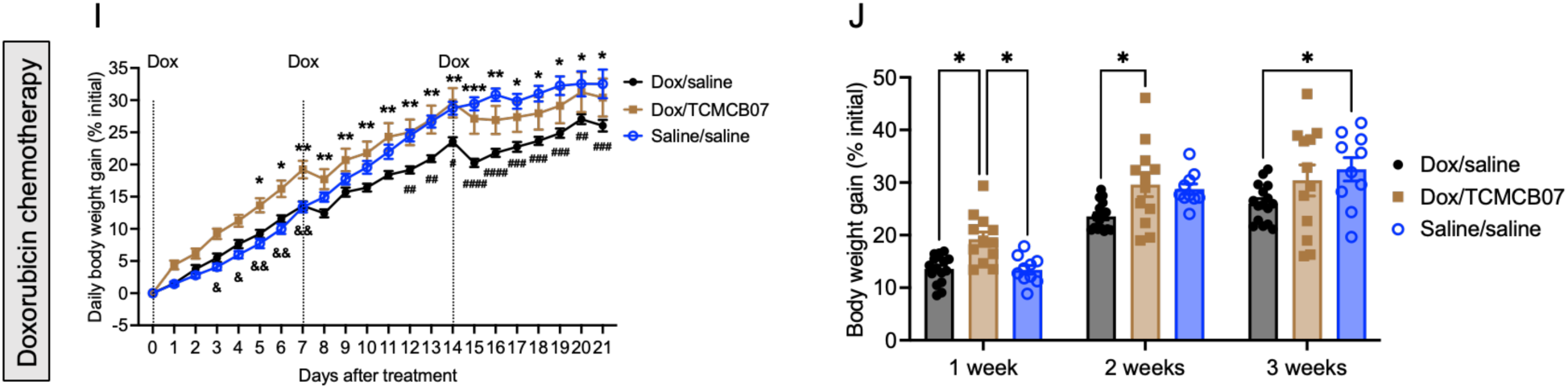
**TCMCB07 treatment preserves body mass throughout multiple cycles of chemotherapy. (A-J**) Daily and weekly body weight gain (% initial body weight) after chemotherapy and TCMCB07 treatment. (**A**) and (**B**) Cisplatin chemotherapy. (**C**) and (**D**) 5-Fluorouracil chemotherapy. (**E**) and (**F**) Cyclophosphamide chemotherapy. (**G**) and (**H**) Vincristine chemotherapy. (**I**) and (**J**) Doxorubicin chemotherapy. All data in (**A**), (**C**), (**E**), (**G**), (**I**) are expressed as mean ± SEM for each group, and all data in (**B**), (**D**), (**F**), (**H**), (**J**) are expressed with each dot representing one sample. *n* = 10-12. (**A**), (**C**), (**E**), (**G**), (**I**), *: Chemotherapy/saline vs Chemotherapy/TCMCB07, #: Chemotherapy/saline vs Saline/saline, &: Chemotherapy/TCMCB07 vs Saline/saline. *, #, &, *P* < 0.05; **, ##, &&, *P* < 0.01; ***, ###, &&&, *P* < 0.001; ****, ####, &&&&, *P* < 0.0001. All data in (**A-H**) were analyzed by Two-way ANOVA, and all data in (**I-L**) were analyzed by One-way ANOVA.

### TCMCB07 treatment attenuates chemotherapy-induced tissue wasting

To assess whether TCMCB07 treatment affects whole body fat and lean mass during multiple cycles of chemotherapy, we measured body composition before and after TCMCB07 treatment. While fat mass decreased significantly in rats after three cycles of cisplatin, 5-FU, cyclophosphamide, or vincristine chemotherapy, rats treated with TCMCB07 fully retained their fat mass (Figure 4A, Supplemental Figure 3A). In contrast, TCMCB07 treatment did not significantly mitigate the loss of lean mass (Figure 4B, Supplemental Figure 3B). Moreover, we observed a significant normalization of heart mass in rats receiving cisplatin/TCMCB07 or 5-FU/TCMCB07 treatment compared to those receiving cisplatin/saline or 5-FU/saline treatment (Figure 4C, Supplemental Figure 3C). However, there was no significant increase in gastrocnemius mass in rats treated with chemotherapy/TCMCB07 compared to those treated with chemotherapy/saline (Figure 4D, Supplemental Figure 3D).

**Figure 4:**
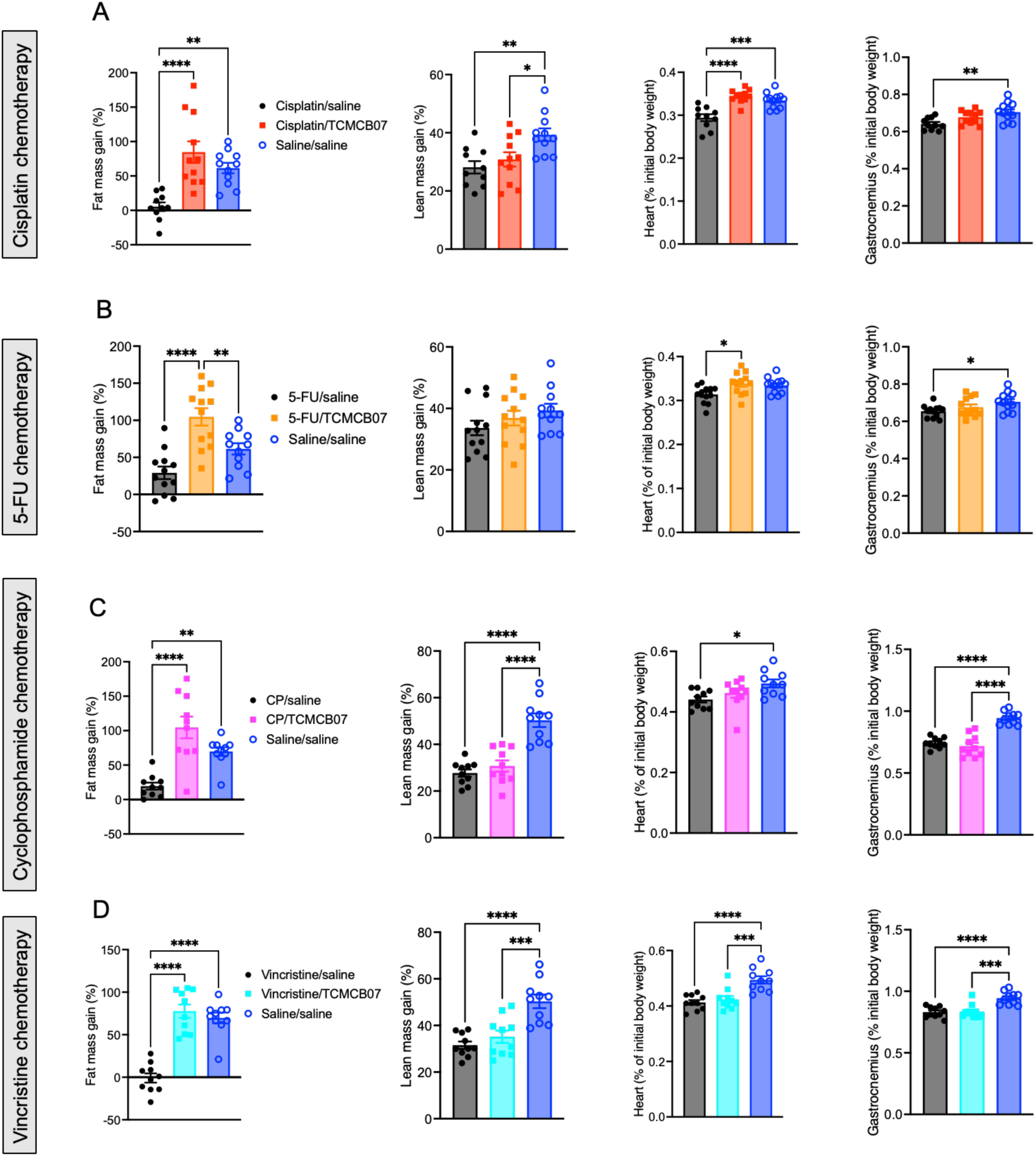
TCMCB07 treatment attenuates chemotherapy-induced tissue wasting. Fat mass gain (% initial), lean mass gain (% initial), heart mass (% initial body weight), and gastrocnemius mass (% initial body weight) after chemotherapy of (**A**) Cisplatin, (**B**) 5-Fluorouracil, (**C**) Cyclophosphamide, (**D**) Vincristine, and TCMCB07 treatment. All data in (**A**-**D**) are expressed with each dot representing one sample. *n* = 10-12. \**P* < 0.05; \*\**P* < 0.01; \*\*\**P* < 0.001; \*\*\*\**P* < 0.0001, One-way ANOVA.

### TCMCB07 + GDF15 antibody combination therapy enhances effectiveness in reversing chemotherapy-induced anorexia and weight loss

Certain chemotherapy agents, particularly cisplatin, raise circulating GDF15 levels (9, 15, 51), which, in turn, suppresses appetite and promotes fat loss and muscle atrophy by activating GFRAL in the brainstem and other GDF15 receptors or pathways (9, 11, 12, 14). We sought to simultaneously target both critical central regulation systems of feeding and metabolism, the hypothalamic melanocortin and GDF15-GFRAL, using a combination therapy involving TCMCB07 and a GDF15 antibody. To test for possible synergistic effects of this combination therapy in reversing anorexia and weight loss following higher doses and multiple cycles of chemotherapy, we first measured GDF15 levels in serum samples collected from TCMCB07/chemotherapy studies. Serum GDF15 levels were elevated in rats receiving chemotherapy with cisplatin, 5-FU, cyclophosphamide, vincristine, or doxorubicin, while no alterations in GDF15 levels were observed with TCMCB07 treatment (Figure 5, A-E). We then challenged rats with a higher dose of cisplatin chemotherapy, followed by administration of the combination therapy of TCMCB07 and GDF15 antibody (Figure 5F). Consistent with previous reports (15), the effectiveness of the GDF15 antibody was validated in mitigating cisplatin-induced anorexia and weight loss. In the present combined treatment study, all three groups of rats received an initial cycle of high-dose (5.0 mg/kg) cisplatin chemotherapy. However, due to mortality observed after the first cycle of chemotherapy, we reduced the dose to 3.0 mg/kg for the second and third cycles (Figure 5G). Of note, either the 5.0 mg/kg or 3.0 mg/kg dosage surpassed the 2.5 mg/kg dose employed in the earlier monotherapy study. Over three cycles of higher-dose cisplatin chemotherapy, rats receiving combination therapy of TCMCB07 + GDF15 antibody exhibited increased daily food intake (Figure 5G), and significant improvement in cumulative (Figure 5H), weekly (Figure 5I), and total food intake (Figure 5J), compared to those receiving IgG or GDF15 antibody monotherapy. Correspondingly, the daily body weight (Supplemental Figure 4A), body weight gain (Figure 6A), and weekly body weight gain (Figure 6B) were significantly higher in rats receiving cisplatin/TCMCB07+GDF15 antibody treatment compared to those receiving cisplatin/saline+IgG or cisplatin/saline+GDF15 treatment. Furthermore, TCMCB07+GDF15 treatment normalized fat mass (Figure 6C, Supplemental Figure 4B), and slightly increased lean mass (Figure 6D, Supplemental Figure 4C) and heart mass (Figure 6E, Supplemental Figure 4D), and gastrocnemius mass (Figure 6F, Supplemental Figure 4E).

**Figure 5:**
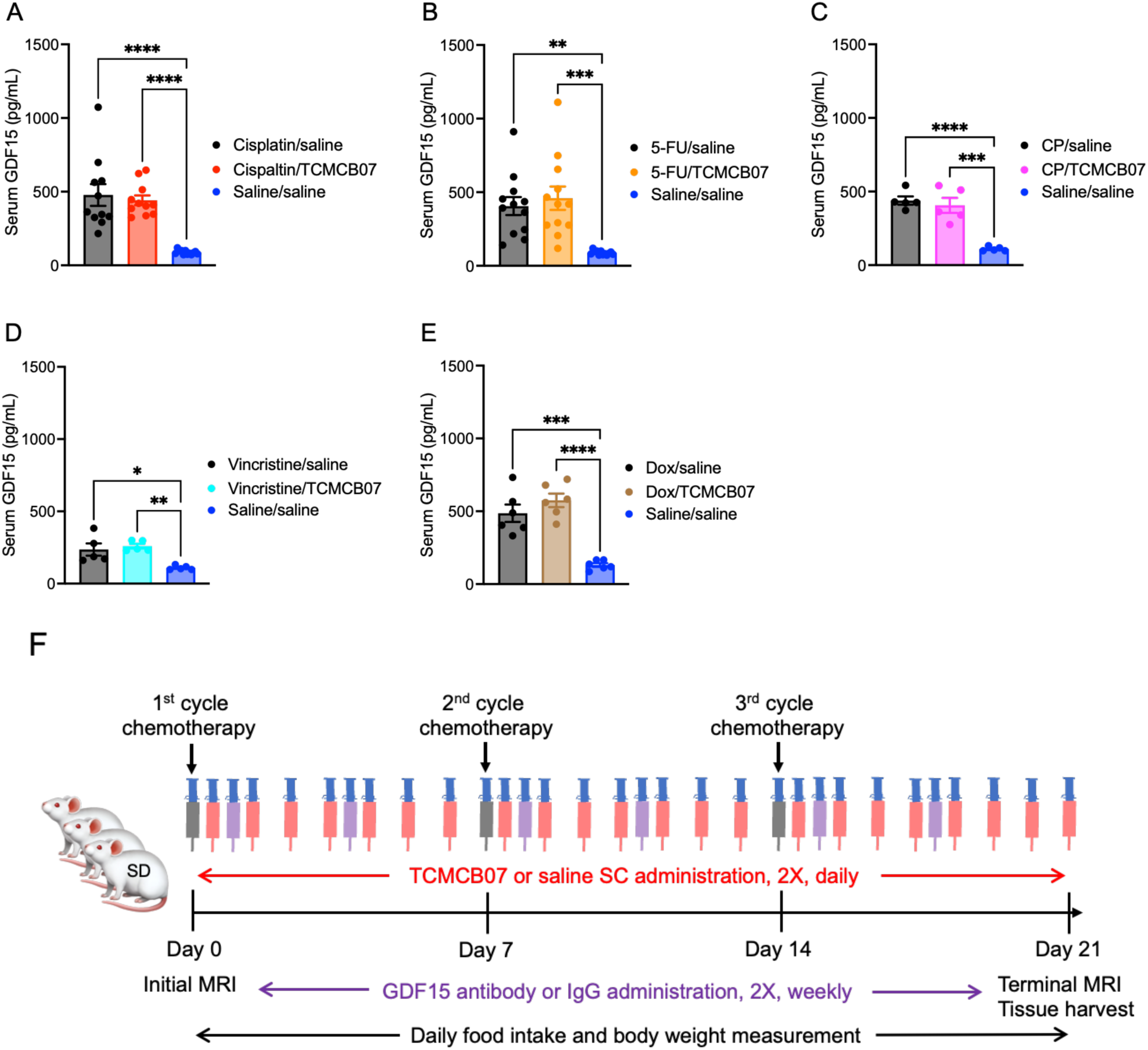
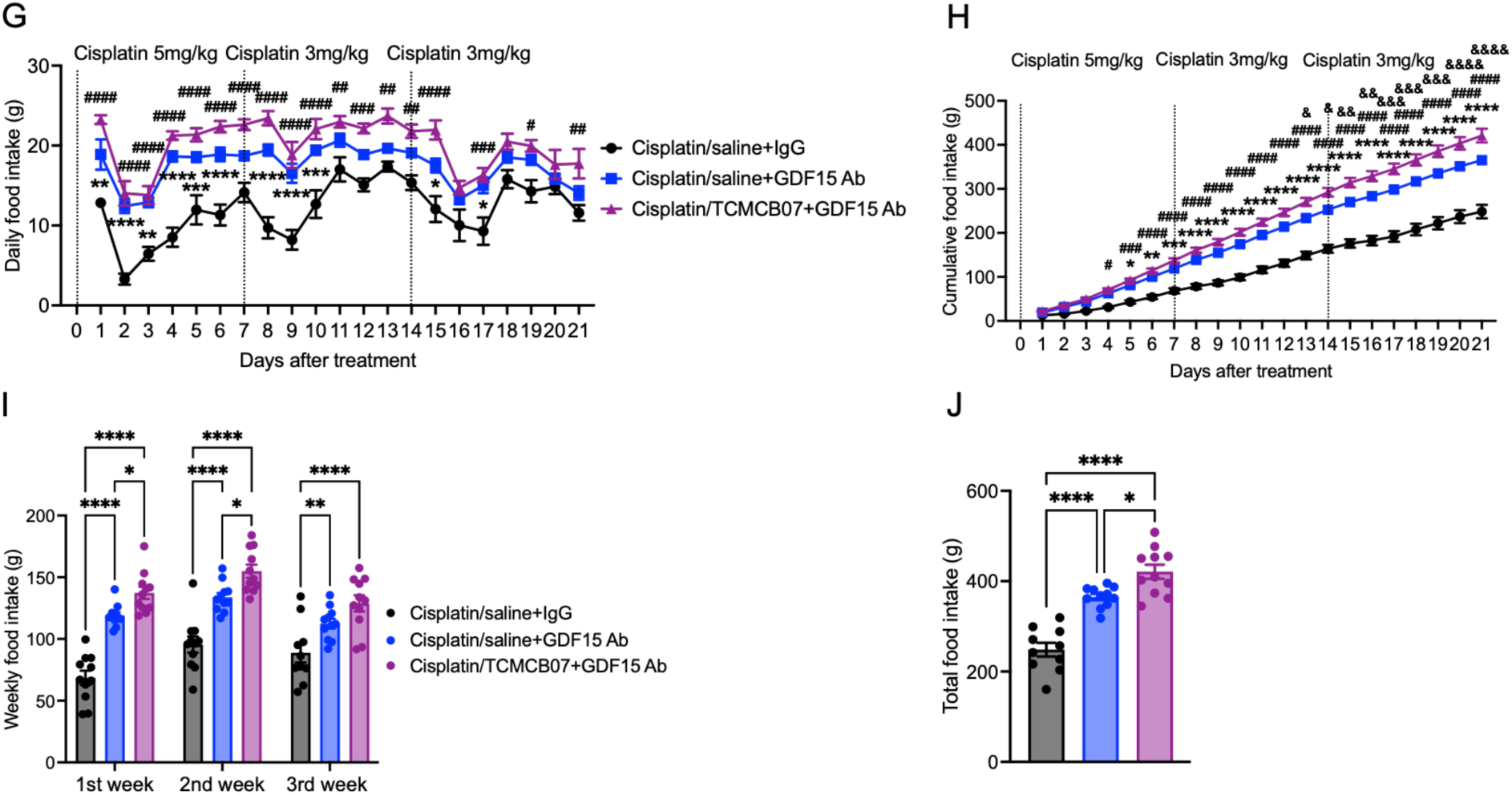
TCMCB07 + GDF15 antibody combination therapy enhances effectiveness in reversing chemotherapy-induced anorexia. **(A**-**E)** SD male rat serum GDF15 concentrations measured from experiments: Cisplatin/TCMCB07 (**A**), 5-Fluorouracil /TCMCB07 (**B**), Cyclophosphamide/TCMCB07 (**C**), Vincristine/TCMCB07 (**D**), and Doxorubicin/TCMCB07 (**E**). (**F**) Schematic of experimental design. All SD male rats were treated with cisplatin chemotherapy via IP injection at a dose of 5.0 mg/kg (1^st^ cycle) or 3.0 mg/kg (2^nd^ and 3^rd^ cycles) once per week for three cycles. All the rats received SC injections twice (2x) daily with either saline or TCMCB07 (3 mg/kg/day) from day 0 to 21. Additionally, all the rats received SC injections twice (2x) weekly with either IgG or GDF15 antibody from day 0 to 21. The dose of TCMCB07 was 3 mg/kg/day, and the dose of GDF15 antibody was 10 mg/kg. Initial and terminal body composition was measured using MRI before and after treatments. Food intake and body weight were monitored daily throughout entire experimental period (day 0-21). At the end of the experiment, tissues were harvested following euthanasia. (**G**) Daily food intake, (**H**) Cumulative food intake, (**I**) Weekly food intake, (**J**) Total food intake after treatment of cisplatin and TCMCB07 in combination with GDF15 antibody. All data in (**A**-**E**), (**I**), and (**J**) are expressed with each dot representing one sample, and all data in (**G**) and (**H**) are expressed as mean ± SEM for each group. (**A**) and (**B**), *n* = 11-12. (**C**-**E**), *n* = 5-6, as blood samples were collected from half of the animals in these experiments. (**G**-**J**), *n* = 10-11. (**G**) and (**H**), *: Cisplatin/saline+IgG vs Cisplatin/saline+GDF15 antibody, #: Cisplatin/saline+IgG vs Cisplatin/TCMCB07+GDF15 antibody, &: Cisplatin/saline+GDF15 antibody vs Cisplatin/TCMCB07+GDF15 antibody. *, #, &, *P* < 0.05; **, ##, &&, *P* < 0.01; ***, ###, &&&, *P* < 0.001; ****, ####, &&&&, *P* < 0.0001. One-way ANOVA (**A**-**E**), (**I**), and (**J**). Two-way ANOVA

**Figure 6:**
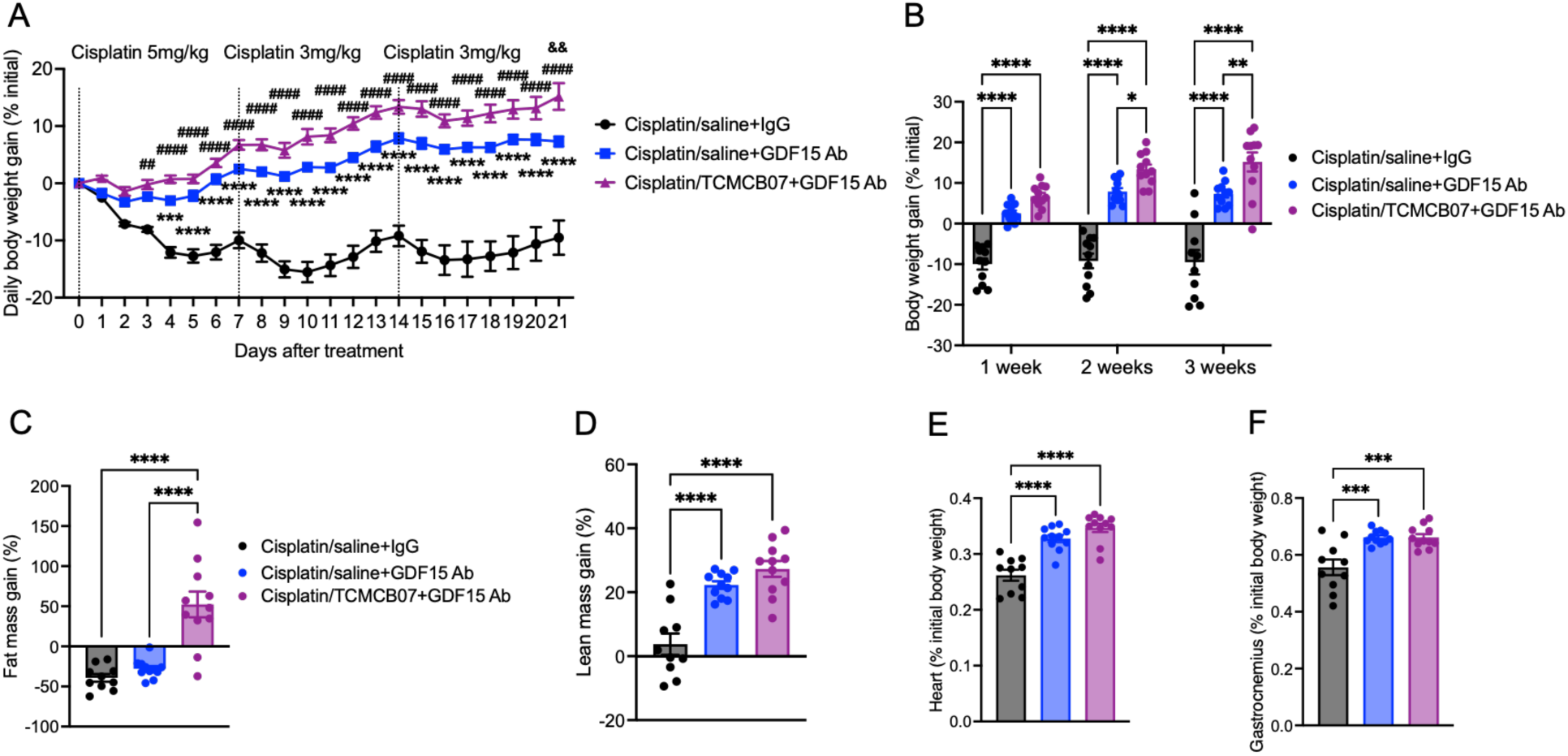
TCMCB07 + GDF15 antibody combination therapy enhances effectiveness in maintaining body and tissue mass during chemotherapy. (**A**) Daily body weight gain (% initial), (**B**) Weekly body weight gain (% initial), (**C**) Fat mass gain (% initial), (**D**) Lean mass gain (% initial), (**E**) Heart mass (% initial body weight), (**F**) Gastrocnemius mass (% initial body weight) after treatment of cisplatin+saline+IgG, cisplatin+saline+GDF15 antibody (Ab), or cisplatin+TCMCB07+GDF15 Ab. All data in (**A**) are expressed as mean ± SEM for each group, and all data in (**B**-**F**) are expressed with each dot representing one sample. *n* = 10-11. (**A**), *: Cisplatin/saline+IgG vs Cisplatin/saline+GDF15 antibody, #: Cisplatin/saline+IgG vs Cisplatin/TCMCB07+GDF15 antibody, &: Cisplatin/saline+GDF15 antibody vs Cisplatin/TCMCB07+GDF15 antibody. \**P* < 0.05; **, ##, &&, *P* < 0.01; ***, *P* < 0.001; ****, ####, *P* < 0.0001. Two-way ANOVA (**A**), One-way ANOVA (**B**-**F**).

### TCMCB07 treatment mitigates anorexia and weight loss in rats with Ward colorectal tumor following combination chemotherapy

To enhance the clinical relevance of this preclinical drug trial, we evaluated TCMCB07’s efficacy in the rat Ward colorectal carcinoma model with combination irinotecan and 5-FU chemotherapy. We first generated the subcutaneous Ward tumor model in Fischer 344 (F344) female rats according to previous studies (37–41), monitoring the tumor growth and tumors’ response to chemotherapy (Figure 7A). We then adapted the doses (50 mg/kg for each agent) and regimen (IP injection, once per week, 5-FU administered 24-h after irinotecan administration) for a total of two cycles of the combination chemotherapy (Figure 7A), previously shown to inhibit the Ward tumor growth and also induce anorexia and weight loss (37–41). Tumor volume was reduced following the first cycle of the chemotherapy, with a nadir on day 4 post-chemotherapy, and then gradually increased until the second chemotherapy treatment (Figure 7B). The second cycle of chemotherapy further suppressed tumor growth (Figure 7B). There was no difference in tumor volume between chemotherapy/TCMCB07 and chemotherapy/saline groups. Chemotherapy induced anorexia and weight loss compared to both baseline levels and the non-tumor (sham-opreated) control group. TCMCB07 treatment mitigated anorexia in tumor/chemotherapy rats compared to saline-treated tumor/chemotherapy rats (Figure 7, C-E). In line with the increased food intake, body weight gain was significantly higher in the tumor/chemotherapy/TCMCB07 group compared to the tumor/chemotherapy/saline group (Figure 7, F and G, Supplemental Figure 6A), This increase in body weight was attributable to a remarkable retention of fat mass and a slight preservation of lean mass (Figure 7, H and I, Supplemental Figure 5, B and C). TCMCB07 treatment also protected cardiac muscle but had no effect on gastrocnemius mass in tumor-bearing rats following the combination chemotherapy (Figure 7, J and K, Supplemental Figure 5, D and E). We did not observe an effect of TCMCB07 treatment on Ward tumor growth, which is consistent with our previous findings from cancer-cachexia studies (31, 34)

**Figure 7:**
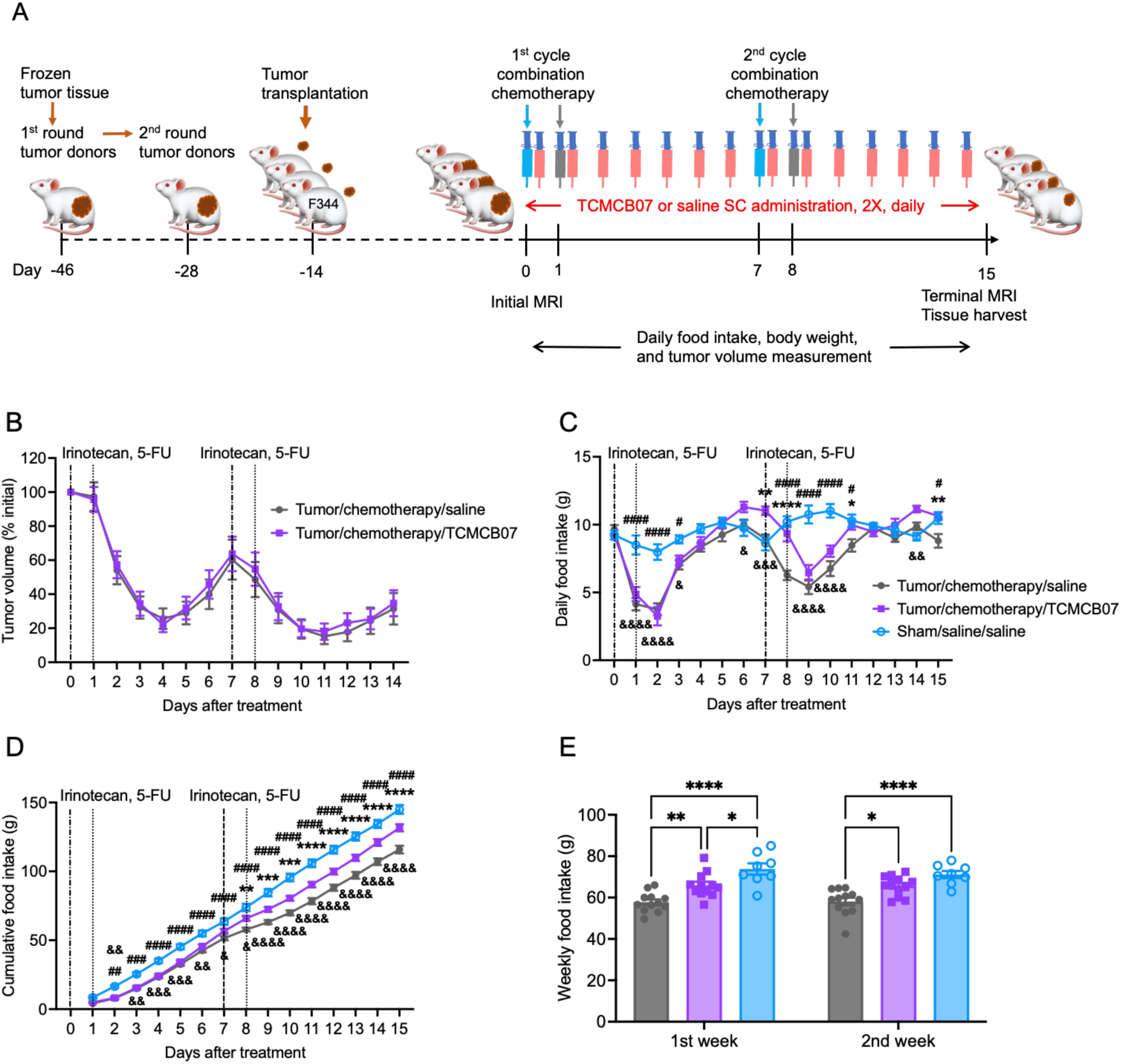
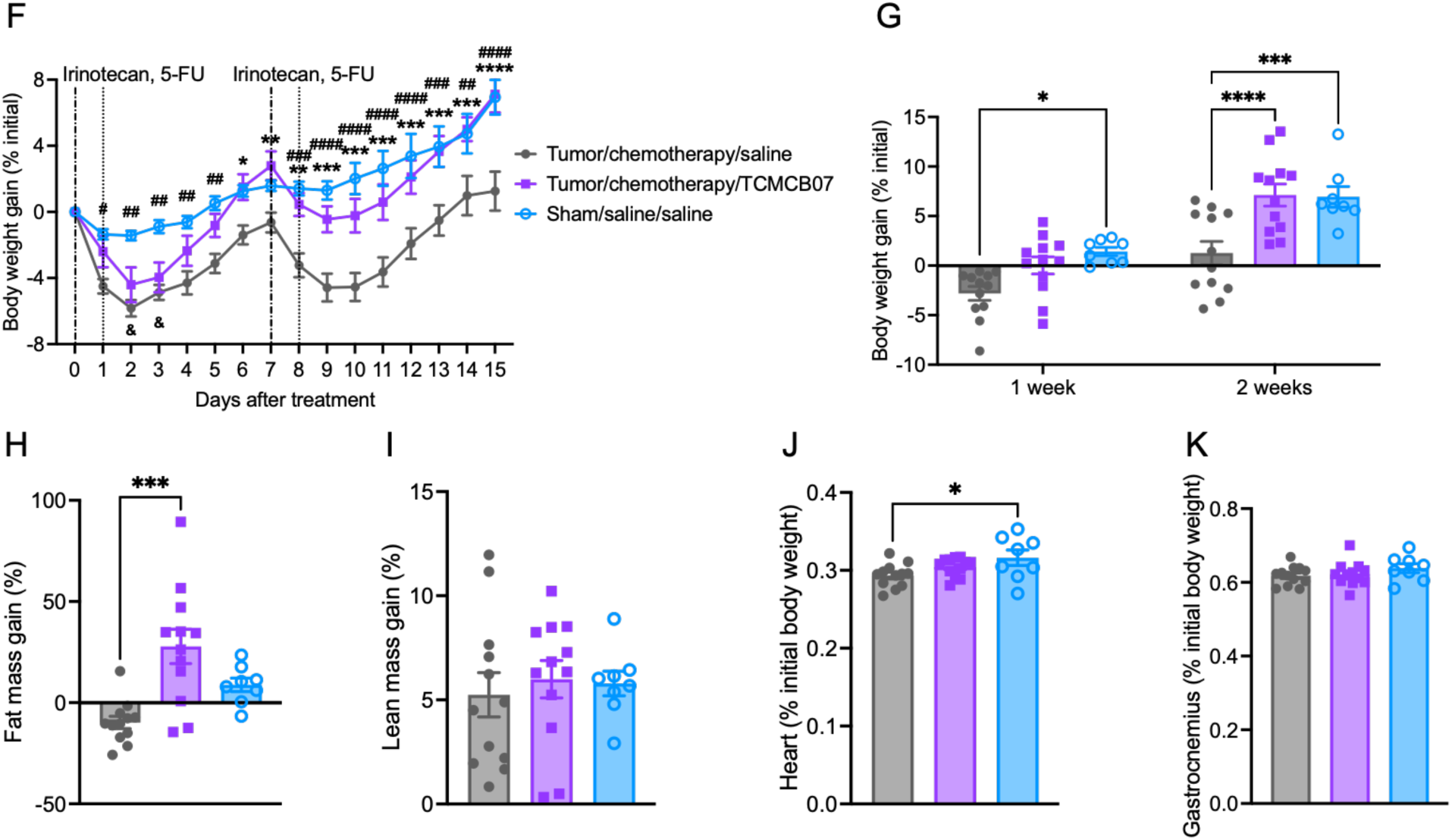
TCMCB07 treatment mitigates anorexia and weight loss in rats with Ward colorectal tumor following combination chemotherapy. (**A**) Experimental design schematic. After passing the frozen tumor tissue through two-rounds of donors, the fresh tumor tissue was subcutaneously implanted into Fischer (F344) female rats. Two weeks later, all rats received combination chemotherapy via IP injection once per week for two cycles: irinotecan (50 mg/kg) on day 0 and day 7, and 5-FU (50 mg/kg) on day 1 and day 8. Additionally, all rats received SC injections twice (2x) daily with either saline or TCMCB07 at a dose of 3 mg/kg from day 0 to day 14. Initial and terminal body composition was measured using MRI prior to and post treatments. Food intake, body weight, and tumor volume were monitored daily from day 0 to day 15. At the end of the experiment, tissues were harvested post-euthanasia. After either chemotherapy or saline and either TCMCB07 or saline treatment, (**B**) Tumor volume change, (**C**) Daily food intake, (**D**) Cumulative food intake, (**E**) Weekly food intake, (**F**) Daily body weight gain (% initial), (**G**) Weekly body weight gain (% initial), (**H**) fat mass gain (% initial), (**I**) Lean mass gain (% initial), (**J**) Heart mass (% initial body weight), (**K**) Gastrocnemius mass (% initial body weight). All data in (**B**-**D**) and (**F**) are expressed as mean ± SEM for each group, and all data in (**E**) and (**G**-**K**) are expressed with each dot representing one sample, *n* = 8-12. (**C**), (**D**) and (**F**), *: Tumor/chemotherapy/saline vs Tumor/chemotherapy/TCMCB07, #: Tumor/chemotherapy/saline vs Sham/saline/saline, &: Tumor/chemotherapy/TCMCB07 vs Sham/saline/saline. *, #, &, *P* < 0.05; **, ##, &&, *P* < 0.01; ***, ###, &&&, *P* < 0.001; ****, ####, &&&&, *P* < 0.0001. Two-way ANOVA (**B***-***D**) and (**F**), One-way ANOVA (**E**) and (**G**-**K**).

### TCMCB07 is detectable in the circulation without causing adverse effects

To ascertain the detectability of administered TCMCB07 in the circulation and assess its correlation with chemotherapy, we quantified TCMCB07 concentrations in rat serum using LC-MS/MS & LC-MRM at the end of both the 21-day cisplatin/TCMCB07 study and a 21-day TCMCB07 test. TCMCB07 was detectable within 0.5-2.5 hours after the last TCMCB07 SC injection (Table 1). Serum TCMCB07 concentrations were higher in rats undergoing chemotherapy (Table 1), suggesting either reduced drug metabolism or reduced clearance due to chemotherapy-induced organ damage. Additionally, hematological parameters in blood samples were analyzed at the end of each study. We observed expected reductions in total leukocyte counts and lymphocyte counts in rats treated with chemotherapy, whereas no significant changes were noted from TCMCB07 treatment (Supplemental Figure 5, A-F). Importantly, in our daily observation of both behavioral responses and overall health status across all groups receiving TCMCB07 treatment, no significant adverse effects or morbidity were attributed to TCMCB07 administration in conjunction with six different chemotherapies and the GDF15 antibody.

**Table 1.** TCMCB07 doses and serum concentrations in rats treated with cisplatin chemotherapy. At the end of 21-day experiments, serum was collected after rats were euthanized. Serum TCMCB07 concentrations were quantitated. Data for the serum concentrations are expressed as mean ± SEM for each group. ***, *p* < 0.001 versus saline/TCMCB07 group.

| Group<br>(n = 10-11) | TCMCB07 doses<br>(mg/kg/day, SC injection) | TCMCB07 serum<br>concentrations (µg/mL) |
| --- | --- | --- |
| Saline/saline | 0.0 | 0.00 |
| Saline/TCMCB07 | 3.0 | 1.48 ± 0.16 |
| Cisplatin/saline | 0.0 | 0.00 |
| Cisplatin/TCMCB07 | 3.0 | 3.34 ± 0.37*** |

## Discussion

In this investigation, to evaluate the broad effectiveness of TCMCB07 in alleviating chemotherapy-induced anorexia and weight loss, we established rat models using six classical agents that represent distinct classes of chemotherapy agents frequently prescribed for various cancer types. While these six chemotherapy agents are commonly employed for cancer treatment, they cause a variety of common side effects and can lead to major organ damage. Significant progress has been made in developing interventions to mitigate these side effects and address organ toxicity (52). However, only one FDA-approved drug, olanzapine, has been shown to improve weight loss in the context of chemotherapy in a randomized trial, and the contexts in which it is effective remain poorly defined (53).

A wide range of chemotherapy dosages and regimens are documented in both literature and clinical practice (43) (54). In preclinical research, the strategies for utilizing chemotherapy in experimental animals predominately depend on the specific purposes of the studies (42, 46–50, 55). However, due to the potency and toxicity, numerous aspects related to animal models of chemotherapy require careful consideration to strike a balance between effectiveness and toxicity. While clinical studies often allow for further dose escalation with extensive supportive care measures such as intravenous hydration, anti-emetics, antihistamines, and corticosteroids, these measures are typically absent in animal studies (43). Moreover, animal species, strains (56), sex, age, body weight, and growth period, can significantly influence responses, effectiveness, and tolerability, thereby markedly impacting investigation outcomes. Additionally, the effects of chemotherapy are not solely determined by the dosage administered in a study but can also be affected by the quality or even the manufacturer of the agents used. For example, we observed variable degrees of sickness responses induced by different suppliers of cisplatin. Furthermore, the dosing regimen is crucial, particularly in multiple cycles of treatment (43). During the initial design of our study, we integrated previous reports and then conducted a series of dose-response experiments in both mouse and rat species. Consistent with existing literature (43), we noted a higher tolerance to the agents we used in this study in mice compared to rats. This difference is likely due to species variations, but more importantly, the total amount of agents administered to rats is at least 10-fold higher compared to mice, as the total amount of drug in preclinical studies is generally calculated based on body weight rather than body surface area. We observed differences even within the same strain of rats: heavier rats exhibited a more pronounced response than lighter ones when given the same dose (in mg/kg). We conclude that precise dosing is critical for the analysis of sickness responses to chemotherapy, and reinforce the importance of dose finding studies, consistent agent quality, and accurate administration volume to achieve this goal. Notably, the doses we selected for our rat model are similar to those used in patients. The clinical doses, according to a prior comparative medicine (43), converted from mg/m^2^ to mg/kg with repeated cycles, are as follows: Cisplatin 2.5 mg/kg, 5-FU 71 mg/kg, Cyclophosphamide 60 mg/kg, Doxorubicin 1.9 mg/kg, and Irinotecan 8.9 mg/kg. In our rat model with multiple cycles, we used the following doses: Cisplatin 2.5, 3.0, 5.0 mg/kg, 5-FU 70 mg/kg, Cyclophosphamide 65 mg/kg, Doxorubicin 2.0 mg/kg, Irinotecan 50 mg/kg.

Our assessment of the effectiveness of TCMCB07 in reversing chemotherapy-induced adverse effects, including anorexia and weight loss, was based on our understanding of underlying mechanisms and previous studies (19, 25, 28, 30, 34, 57). Encouragingly, our data demonstrated the promising potential of TCMCB07 treatment to provide beneficial effects on both food intake and body weight in rats treated with a variety of chemotherapeutics for multiple cycles. In studies involving cisplatin or 5-FU chemotherapy, TCMCB07 treatment fully restored the loss of appetite and body weight following each chemotherapy cycle, highlighting its potential to prevent chemotherapy dose reductions in a clinical setting. In contrast to the cisplatin and 5-FU studies, TCMCB07 treatment did not fully reverse appetite and weight loss in rats undergoing cyclophosphamide, vincristine, or doxorubicin chemotherapy. This could be attributed to the higher toxicity of these chemotherapy agents at the experimental dosage levels and highlight the need for multiple orexigenic agents to address this heterogeneous toxicity.

Consistent with previous reports (11, 58), tissue wasting was observed following three cycles of chemotherapy of cisplatin, 5-FU, cyclophosphamide, or vincristine. Body composition analyses revealed significantly lower fat and lean mass among rats undergoing chemotherapy, including a dramatic depletion in fat mass. Furthermore, we observed cardiac and skeletal muscle loss following chemotherapy treatment. TCMCB07 treatment protected heart mass following cisplatin or 5-FU, and slightly improved lean mass across all chemotherapy-treated models. This predicts a clinical benefit of melanocortin antagonism in patients receiving these agents and provides a rationale for clinical investigation of this therapeutic modality. In contrast, we did not observe a significant protective effect of TCMCB07 treatment on gastrocnemius mass in all individual studies, consistent with others’ observations (15). In our previous cachexia studies with both rat models and dogs (31, 34), TCMCB07 showed muscle improvement, but this was not observed in the chemotherapy study. We attribute this to several factors: a) TCMCB07 does not directly stimulate protein synthesis or promote muscle proliferation and differentiation. Instead, it indirectly aids in muscle preservation by enhancing nutrient availability and general anabolism. b) Chemotherapy-induced muscle loss occurs due to a reduction in food intake, toxicity-related tissue damage (59, 60), and inhibition of protein synthesis in skeletal muscle (61). c) The molecular mechanisms of muscle loss induced by chemotherapy may differ from those involved in cancer cachexia-associated muscle wasting (62–65). Therefore, despite adequate nutritional availability, the anabolic potential of chemotherapy-treated skeletal muscle may be impaired.

To enhance the effectiveness and outcomes of interventions targeting the reversal of chemotherapy-induced side effects, it will be inevitable to develop novel treatment strategies that integrate various therapeutic approaches aimed at achieving synergistic effects. Accumulating evidence demonstrates that GDF15 is a key negative mediator of appetite and body weight via its action on the GFRAL receptor and potentially other pathways and receptors. Moreover, circulating GDF15 levels are elevated following chemotherapy with specific agents, such as cisplatin, and beneficial effects were observed by neutralizing the increased GDF15 (15). In our study, chemotherapy with five agents at the administered doses and regimens resulted in a similar elevation of circulating GDF15. It is reasonable to propose that combining both TCMCB07 and a GDF15 antibody would enhance therapeutic effectiveness. While TCMCB07 stimulates feeding and anabolism via the central melanocortin system in the forebrain, the GDF15 antibody attenuates GDF15-GFRAL-mediated feeding suppression in the hindbrain. Using a specific GDF15 antibody, we observed an augmented effectiveness from the combination therapy in rats treated with a higher dose of cisplatin, as evidenced by improved food intake and body weight gain. Additionally, the combined treatment exhibited a more pronounced preservation in fat mass and a positive trend towards an increase in lean mass and heart mass. To our knowledge, this is the first instance of this combinatorial approach being applied to alleviating anorexia and weight loss induced by chemotherapy, supporting the potential for TCMCB07 treatment to be paired with other therapies.

It is essential to assess TCMCB07’S efficacy both with chemotherapy alone and in combination with cancer and chemotherapy. The former context reflects a common use of chemotherapy in the adjuvant setting, when there is no evidence of active disease. Furthermore, testing the effects alongside a single chemotherapy agent in healthy animals is important to identify the specific impact without potential interference from other variables. To better understand the clinical potential of TCMCB07, it is equally important to validate whether these effects persist or change in the presence of cancer and combined chemotherapy regimens (63). In clinical practice, multiple chemotherapy agents are commonly used to treat patients with tumors, both in the palliative and neoadjuvant settings. To mimic this, we utilized the Ward colorectal tumor model in female F344 rats and employed a combination chemotherapy regimen of irinotecan and 5-FU. Notably, female F344 rats exhibit absolute weight loss following chemotherapy due to their slower growth and weight gain compared to SD rats (66–68). As previously reported, the combination chemotherapy of irinotecan and 5-FU effectively reduced tumor size in this model. This combination, previously used in multiple rat studies, closely resembles FOLFIRI, a regimen commonly used to treat colorectal cancer (37–40). TCMCB07 administration alleviated anorexia and weight loss in the chemotherapy-treated tumor-bearing rats. Since TCMCB07 has previously shown efficacy in ameliorating cancer cachexia in both rats and dogs (31, 34), it is rational to speculate that TCMCB07 treatment will be beneficial for cancer patients by mitigating both cancer- and chemotherapy-induced anorexia and weight loss.

Given that MC4R antagonists target the central melanocortin system, the ability of a drug to penetrate the blood-brain barrier (BBB) is essential for its efficacy. To develop effective drugs in this class, one of the primary challenges is that many peptides exhibit positive effects only with direct central delivery, such as AgRP and SHU9119 (melanocortin antagonists), or melanocortin-II (melanocortin agonist), but have no effects when administered peripherally (69–71). While TCMCB07 levels in the central nervous system were not measured in this study, its consistent ability to stimulate feeding and attenuate weight loss through peripheral treatment strongly implies direct activity within the brain. We found that serum TCMCB07 was detectable within 0.5-2.5 hours after final dosing, and that serum concentration correlated with the duration of exposure. Notably, we observed significantly higher concentrations of TCMCB07 in rats receiving chemotherapy, indicating decreased drug metabolism or clearance (42, 46).

Apart from chemotherapy-induced toxicity, no additional adverse effects were observed during TCMCB07 monotherapy or combination therapy with a GDF15 antibody in rats undergoing chemotherapy. While we previously conducted a series of standard evaluations for TCMCB07’s safety in various species, including rats, dogs, and humans (31, 32, 34–36), this study marks the first time we have tested the drug candidate in combination with chemotherapy agents. Encouragingly, we observed no increased toxicity when TCMCB07 was combined with other chemotherapy agents even during a prolonged administration period of 21 days.

This study had several key limitations. Firstly, we predominantly used SD rats in our experiments, chosen for its widespread use in various research fields, easy availability, and cost-effectiveness compared to inbred strains (66). These characteristics made SD rats a suitable choice for our studies, which were conducted across multiple locations in both the USA and China. However, like other common outbred strains such as Wistar, SD rats naturally exhibit continuous weight gain during the ages used in our studies (66–68). Male SD rats, even at 15 weeks old and weighing over 500 grams, continue to gain weight rapidly. The rapid and continuous weight gain in SD rats presented a challenge, as chemotherapy inhibited weight gain instead of inducing absolute weight loss. Although significant relative weight loss was observed compared to non-chemotherapy-treated animals in all related experiments, this limitation introduces some confusion regarding translational relevance and clinical implications. To address this, we used an inbred strain— female F344 rats—and further validated the effects of TCMCB07 following tumor growth and combination chemotherapy. Due to their slower growth rate, F344 rats allowed us to model absolute weight loss relative to baseline following chemotherapy and confirmed the ability of TCMCB07 to reverse this weight loss. Second, although IP injection is the most commonly used method in rodent chemotherapy models, the intravenous (IV) route better reflects clinical practice. However, we chose IP injection due to the considerable challenges associated with IV administration in rats. Unlike in clinical settings, IV injection in rats is technically challenging and can cause significant stress-induced changes in feeding behavior. While our models effectively replicated chemotherapy-induced side effects such as anorexia and weight loss, we recognize that IP injection may result in different pharmacokinetics compared to IV injection (63), potentially affecting the animals’ responses and the interactions between chemotherapy and TCMCB07 treatment. Third, selecting doses of the six chemotheraputic agents that reflect clinical doses posed a significant challenge. While it is important to consider clinical doses (converted from mg/m^2^ to mg/kg for animals) to enhance translational relevance and clinical implications, the biological differences between humans and rats are equally critical. Rats are not simply miniature versions of humans (67). Although the doses of each single agent used in this study were similar to the clinical doses (43), confirming whether these doses are equivalent or substantially different from those used in clinical settings is difficult due to differences in tolerance and metabolism between species. We instead relied upon titrating dose to the desired adverse effects—anorexia and weight loss—which we feel provides a more relevant translational model. Further studies in humans are required to ascertain whether the beneficial effects we observed in these preclinical studies will translate to patients.

## Conclusion

In conclusion, this preclinical study demonstrates that peripheral administration of TCMCB07 increases food intake, mitigates weight loss, and significantly reduces tissue wasting over multiple cycles of various chemotherapy regimens, including combination chemotherapy in a tumor model that mirrors clinical scenarios. Moreover, the combination of TCMCB07 with a GDF15 antibody enhances treatment outcomes. These findings highlight TCMCB07 as a promising drug candidate with strong potential to alleviate chemotherapy-induced anorexia and weight loss, aligning with previous studies on its effectiveness in treating cachexia. Additionally, this study provides preliminary evidence supporting the potential of TCMCB07, in combination with other drugs, to effectively combat severe anorexia and weight loss induced by chemotherapy. TCMCB07 is expected to benefit many cancer patients undergoing chemotherapy.

## Methods

### Rats

Sprague-Dawley (SD) male rats at age of 8 week weighing 200-225 g and Fischer 344 (F344) female rats at age of 12 week weighing 130-140 g were obtained from Charles River Laboratories (Wilmington, MA, USA) or Beijing Vital River Laboratory Technology (Beijing, China), and housed in animal facilities with controlled conditions, including a temperature of 20-22 ^0^C and a 12-h light-dark cycle. The rats had *ad libitum* access to water and food (Purina rodent diet 5001; Purina Mills, St. Louis, MO, USA). After being individually housed for at least 7 days for acclimation, before experiments, rats were assigned into treatment and control groups using body weight (day 0) to counterbalance groups. Rat receiving chemotherapy were monitored closely and euthanized when reaching the endpoints set by the chemotherapy study policy. Our study examined both male and female rats to test whether there was sexual dimorphism in response to chemotherapies and TCMCB07 treatment. All experiments were conducted following the guidelines of the National Institutes of Health Guide for the Care and Use of Laboratory Animals and were approved by the Institutional Animals Care and Use Committee of Oregon Health & Science University and WuXi IACUC standard animal procedures (IACUC protocol number GP02-QD009-2022v1.0.).

### TCMCB07 compound and administration

TCMCB07 was designed and provided by Endevica Bio (Northbrook, IL, USA). According to previous observations, an effective dose of TCMCB07 at 3 mg/kg/day was chosen, and administration route was subcutaneous (SC) injection, to avoid first pass metabolism. To maintain the circulating concentration, one dose of TCMCB07 was split into two SC injections (1.5 mg/kg x two injections) performed in morning (9-10am) and evening (5-6pm). Control animals received an equivalent volume of saline SC injections.

### Chemotherapy agents and regimen

The following six medical grade chemotherapy agents were used: cisplatin (Teva, North Wales, PA), 5-Fluorouracil (Xiromed, LLC, Florham Park, NJ, India), cyclophosphamide (Jiangsu Hengrui Pharmaceuticals, China), vincristine (Shenzhen Main Luck Pharmaceuticals, China), doxorubicin (ShanXi PUDE Pharmaceuticals, China), and irinotecan (Pfizer, PGS Pearl River, NY, USA). The concentrations of the agents were as follows: cisplatin 1 mg/mL, 5-Fluorouracil 50 mg/mL, and irinotecan 20 mg/mL. Upon reconstitution, the concentrations of the agents were: cyclophosphamide 20 mg/mL (in saline), vincristine 0.2 mg/mL (in PBS), and doxorubicin 2 mg/mL (in saline). Atropine (1 mg/kg, subcutaneous) was administered immediately prior to each irinotecan injection to alleviate early onset cholinergic symptoms (72). The doses of chemotherapy agents were selected through a series of dose-response experiments aiming to induce 10 to 30% weight loss compared to rats not receiving chemotherapy (15). In addition, these selected doses did not result in severe morbidity or mortality based on the observations from our doses-response experiments and the literature. The selected doses were as follows (in mg/kg): cisplatin 2.5, 3.0, or 5.0 (with the 5.0 mg/kg dose given only during the first cycle), 5-FU 70 or 50 (with the dose of 50 mg/kg administered in the combination with irinotecan), cyclophosphamide 65, vincristine 0.27, doxorubicin 2.0, and irinotecan 50 mg/kg in the combination with 5-FU. All chemotherapy agents were accurately administered via IP injection using insulin syringes, once per week for a total of three cycles of a single chemotherapy or two cycles of combination chemotherapy. For the combination chemotherapy, irinotecan was administered 24 h before 5-Fluorouracil administration (39). Control animals received an equivalent volume of saline via IP injections. Euthanasia criteria were adhered to as per the IACUC study protocol. Data collected from animals reaching euthanasia criteria before the designed experimental endpoint were excluded from statistical analysis.

### GDF15 antibody and administration

Anti-GDF15 antibody was manufactured by Sanyou Bio (Shanghai, China), based on the published protein sequence of the mAB1 GDF15 antibody, GDF15 antibody (GDF15-001, Ponsegromab, Pfizer). The control antibody (rat IgG2a) was also produced by Sanyou Bio. The concentrations of both anti-GDF15 antibody and IgG2a control were verified by the manufacturer. The anti-GDF15 antibody or IgG2a was administered at dose of 10 mg/kg via SC injection, twice per week for a total of three weeks. The dose was determined based on previous reports (15). Control animals received an equivalent volume of saline via SC injection.

### Ward tumor model

Rat Ward colorectal carcinoma tumor model was generated as described in previous studies (37–41). Briefly, the frozen Ward tumor tissue was subcutaneously implanted into the first round of F344 rat donors. 18 days later, the fresh tumor tissue from the donors was transplanted into the second round of donors to grow tumors for 14 days. Fresh tumor fragments (0.05 g) were subcutaneously implanted into the right flank of experimental F344 rats via a trocar under slight isoflurane anesthesia. Subcutaneous implantation was chosen to facilitate continuous evaluation of tumor growth and response to the chemotherapy. Rats in sham control group recived subcutaneous PBS injection. Tumor volume was monitored every other day prior to chemotherapy and daily post-chemotherapy. Tumors were measured in three dimensions with a caliper: the length (L), the width (W), the height (H). The tumor volume was calculated as following: tumor volume (cm^3^) = 0.5 x L (cm) x W (cm) x H (cm) (38, 73). After approximately two weeks of tumor growth, when tumor volume reached ∼2 cm^3^, irinotecan and 5-FU combination chemotherapy and TCMCB07 treatment were initiated. During two cycles of the combination chemotherapy, tumor volume change (%) in each tumor rat was compared to the baseline volume (day 0) (38).

### Food intake and body weight measurement

We measured food intake and body weight daily at the same time of day (3 hours after lights on) from the day of starting treatment (day 0) until the day of termination. Because chemotherapy-related side effects distress animals, some sick rats produced considerable food spillage (orts or crumbs) as the experiments progressed. To ensure accurate measurement of food intake, we accounted for uningested food loss by screening cage bedding and subtracting orts from total food amount reduction. We also monitored the overall health condition of animals daily to ensure no animals were moribund.

### Body composition analysis

Body composition (fat mass and lean mass) was analyzed twice on the day of treatment and the end of study prior to tissue collection via EchoMRI (4-in-1, Live Animal Composition Analyzer; Echo Medical System, Houston, TX, USA).

### Blood and tissue collection

When experimental animals reached terminal time points, we conducted blood and tissue collection immediately after the terminal MRI scan. Animals were deeply anesthetized with isoflurane, and blood was collected via cardiac puncture. Approximate 200 μL of blood was placed into a 1-mL EDTA blood collection tube for hematology assay, and the remaining blood was placed in a 10-mL serum collection tube. Serum was isolated, aliquoted, and stored at −80 ^0^C until analysis. Additionally, we dissected and weighed the heart and gastrocnemii from bilateral hindlimbs. The average of both gastrocnemius masses was used in the final analysis.

### Hematology assay

Whole blood was subjected to analysis using a veterinary hematology analyzer (HemaVet, 950FS, Drew Scientific, Oxford CT, USA) to measure various hematological parameters, including total leucocyte counts, leucocyte differential (neutrophils, lymphocytes, monocytes, eosinophils, and basophils), erythrocytes, hemoglobin concentration, hematocrit, mean corpuscular volume, mean corpuscular hemoglobin, mean corpuscular hemoglobin concentration, and thrombocytes.

### Enzyme-linked immunosorbent assay (ELISA)

Concentrations of GDF-15 in rat serum were determined using enzyme-linked immunosorbent assay (ELISA) kits following the manufacturers’ instructions (R&D Systems, catalog # DY957).

### Quantitation of serum TCMCB07

At the end of the 21-day studies, rats were euthanized and blood was collected within 0.5-2.5 hours after the last TCMCB07 or saline injection. The serum samples were submitted to the Charles W Gehrke Proteomics Center at University of Missouri-Columbia for analysis of TCMCB07 concentration via liquid chromatograph-tandem mass spectrometry with multiple reaction monitoring (LC-MS/MS & LC-MRM). A standard curve was established using the same batch of TCMCB07 used in the animal experiments. Non-treated rat serum samples served as the blank control. A client specific method was developed for TCMCB07 quantitation. The serum TCMCB07 concentrations (µg/mL) was reported.

### Statistical analysis

Statistical analyses were conducted using GraphPad Prism 10.0 software. Quantitative data are reported as mean ± standard error. To compare two groups, a two-tailed unpaired Student t-test was used. When comparing more than two groups, one-way analysis of variance (ANOVA) was utilized. For comparing multiple time points and treatment groups, unless otherwise specified in the figure legends, two-way ANOVA with Tukey’s multiple comparisons test was used. Statistical significance was considered at a p-value < 0.05 for all data analyses. All measurements were taken from distinct samples, ensuring that no duplications occurred from the same samples.

### Study approval

Animal studies were approved by the IACUC of the Oregon Health & Science University and conducted according to the NIH Guide for the Care and Use of Laboratory Animals (National Academies Press, 2011).

## Data availability

All data associated with this study are available in the main text, main figures, and supplemental materials, or supporting data values file. There are no restrictions on data availability. Values for all graphs are reported in the [Supporting Data Values](https://www.jci.org/articles/view/161987#sd) file.

## Author contributions

XZ, DLM, and RP conceived and designed the study. KAG designed TCMCB07. XZ, SN, MAN, PRL, PB, XC, and QG conducted experiments. EZ organized and oversaw some of the experiments conducted in WuXi, China. XZ, PB, and DLM analyzed the data. AJG contributed to discussion and data interpretation. XZ wrote the manuscript with input from the other authors. DLM and AJG reviewed and edited the manuscript. All authors approved the final version of the manuscript.

## Acknowledgments

This work was supported by NIH NCI R01 CA257452 and CA264133 (to DLM), two research grants from Brenden-Colson Center for Pancreatic Care at OHSU (to DLM), and two research grants from Endevica Bio (to DLM). We thank Dr. Vickie Baracos and Abha Dunichand-Hoedl for generously providing the Ward colorectal tumor tissue and protocol. The graphical abstract was generated using BioRender (BioRender.com).

