## Supplemental Figures for "Melanocortin-4 receptor antagonist TCMCB07 alleviates chemotherapy-induced anorexia and weight loss"

### Slide 1
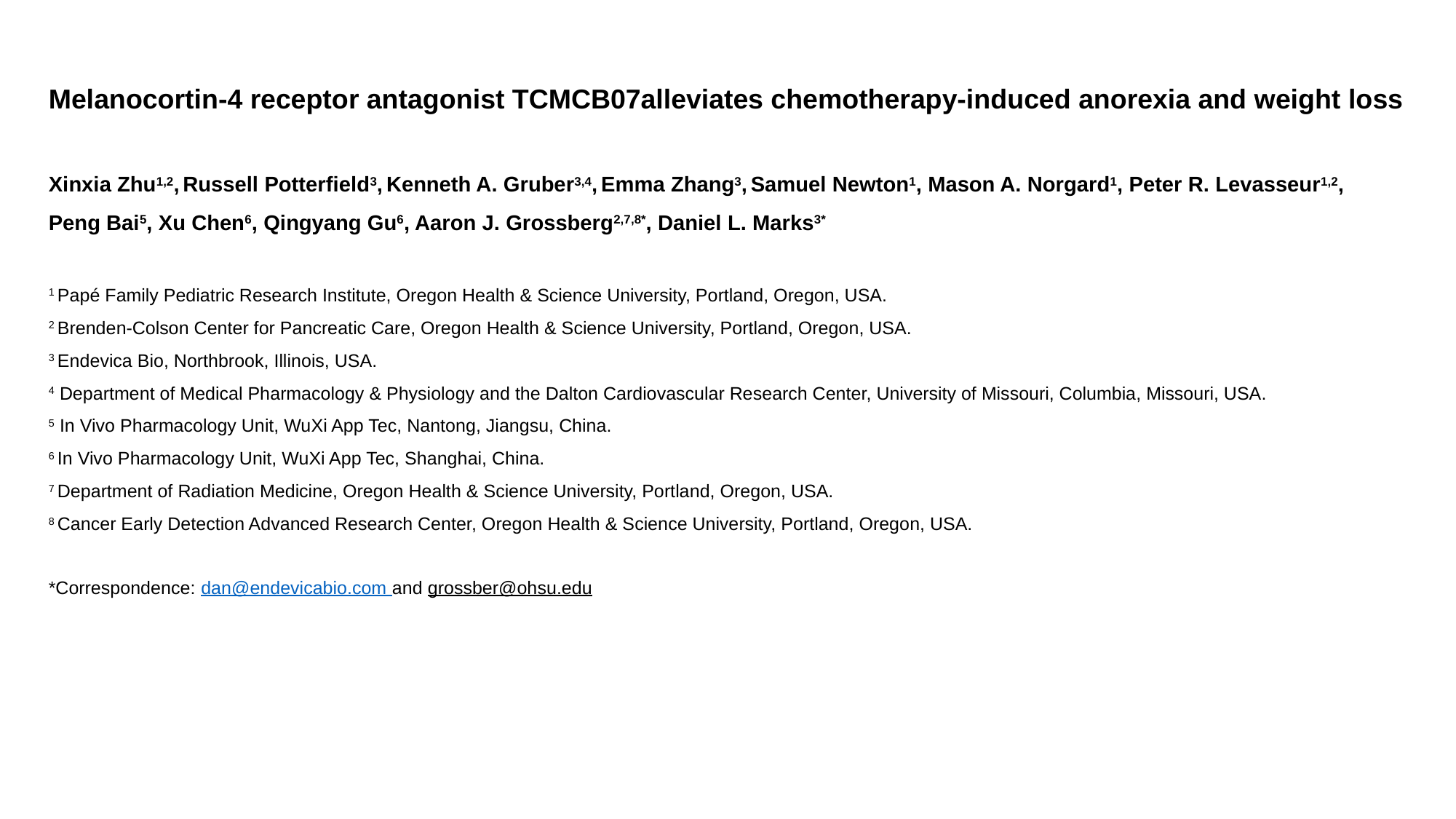

Melanocortin-4 receptor antagonist TCMCB07alleviates chemotherapy-induced anorexia and weight loss
Xinxia Zhu^1,2^, Russell Potterfield^3^, Kenneth A. Gruber^3,4^, Emma Zhang^3^, Samuel Newton^1^, Mason A. Norgard^1^, Peter R. Levasseur^1,2^,
Peng Bai^5^, Xu Chen^6^, Qingyang Gu^6^, Aaron J. Grossberg^2,7,8*^, Daniel L. Marks^3*^
^1^ Papé Family Pediatric Research Institute, Oregon Health & Science University, Portland, Oregon, USA.
^2^ Brenden-Colson Center for Pancreatic Care, Oregon Health & Science University, Portland, Oregon, USA.
^3^ Endevica Bio, Northbrook, Illinois, USA.
^4^ Department of Medical Pharmacology & Physiology and the Dalton Cardiovascular Research Center, University of Missouri, Columbia, Missouri, USA.
^5^ In Vivo Pharmacology Unit, WuXi App Tec, Nantong, Jiangsu, China.
^6^ In Vivo Pharmacology Unit, WuXi App Tec, Shanghai, China.
^7^ Department of Radiation Medicine, Oregon Health & Science University, Portland, Oregon, USA.
^8^ Cancer Early Detection Advanced Research Center, Oregon Health & Science University, Portland, Oregon, USA.

### Slide 2
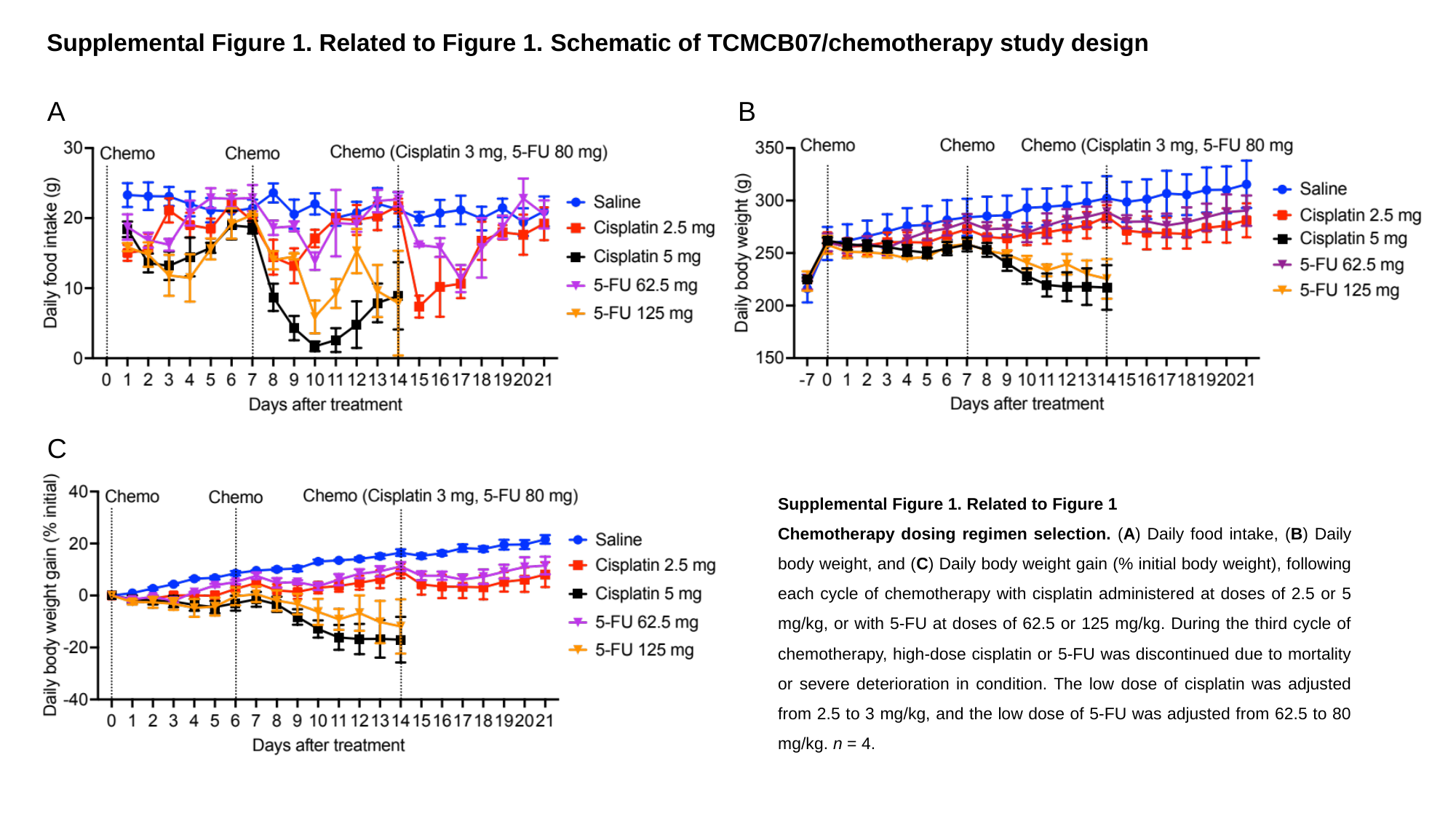

Supplemental Figure 1. Related to Figure 1. Schematic of TCMCB07/chemotherapy study design
A
B
C
Supplemental Figure 1. Related to Figure 1
Chemotherapy dosing regimen selection. (A) Daily food intake, (B) Daily body weight, and (C) Daily body weight gain (% initial body weight), following each cycle of chemotherapy with cisplatin administered at doses of 2.5 or 5 mg/kg, or with 5-FU at doses of 62.5 or 125 mg/kg. During the third cycle of chemotherapy, high-dose cisplatin or 5-FU was discontinued due to mortality or severe deterioration in condition. The low dose of cisplatin was adjusted from 2.5 to 3 mg/kg, and the low dose of 5-FU was adjusted from 62.5 to 80 mg/kg. n = 4.

### Slide 3
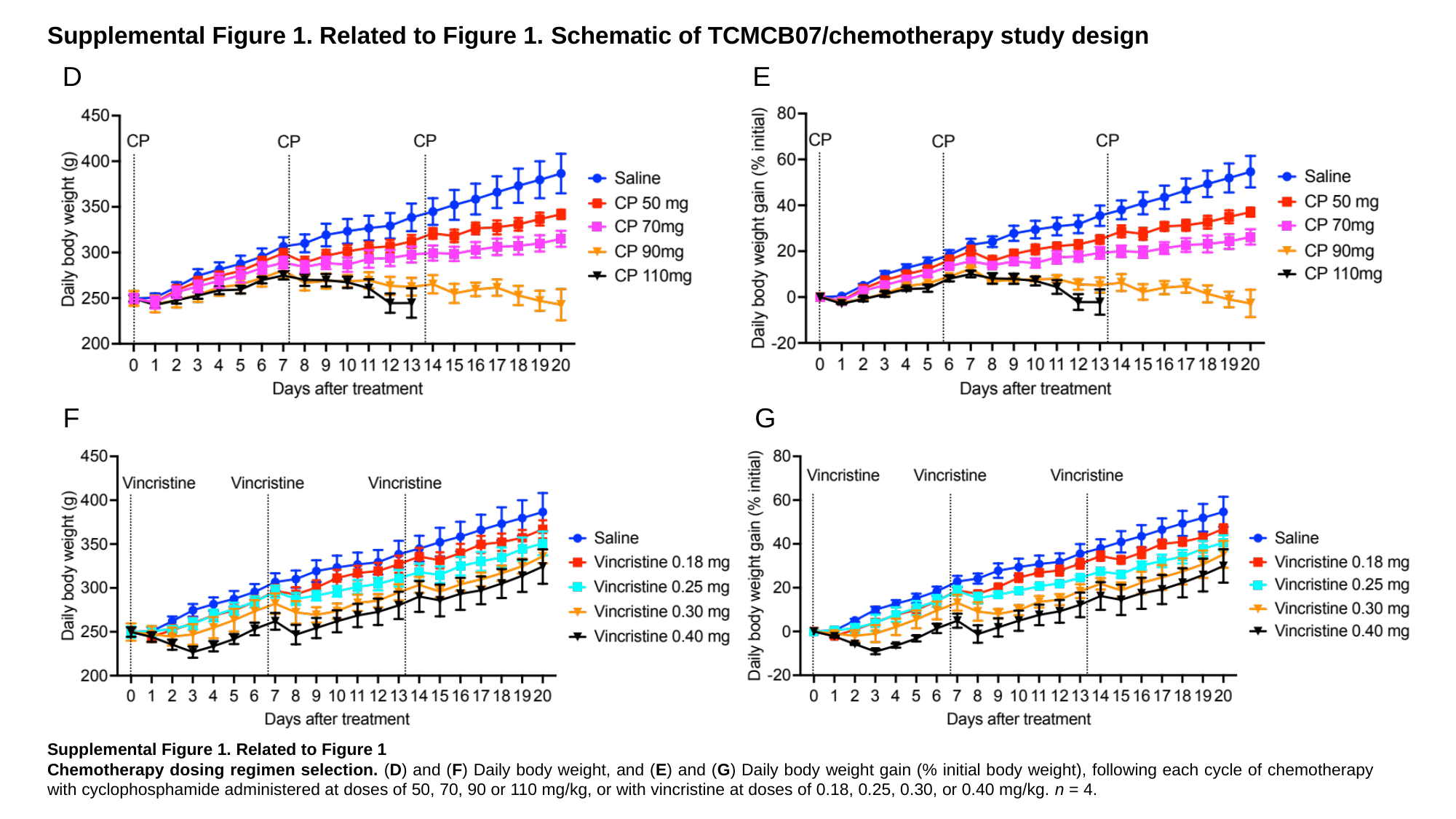

Supplemental Figure 1. Related to Figure 1. Schematic of TCMCB07/chemotherapy study design
D
E
F
G
Supplemental Figure 1. Related to Figure 1
Chemotherapy dosing regimen selection. (D) and (F) Daily body weight, and (E) and (G) Daily body weight gain (% initial body weight), following each cycle of chemotherapy with cyclophosphamide administered at doses of 50, 70, 90 or 110 mg/kg, or with vincristine at doses of 0.18, 0.25, 0.30, or 0.40 mg/kg. n = 4.

### Slide 4
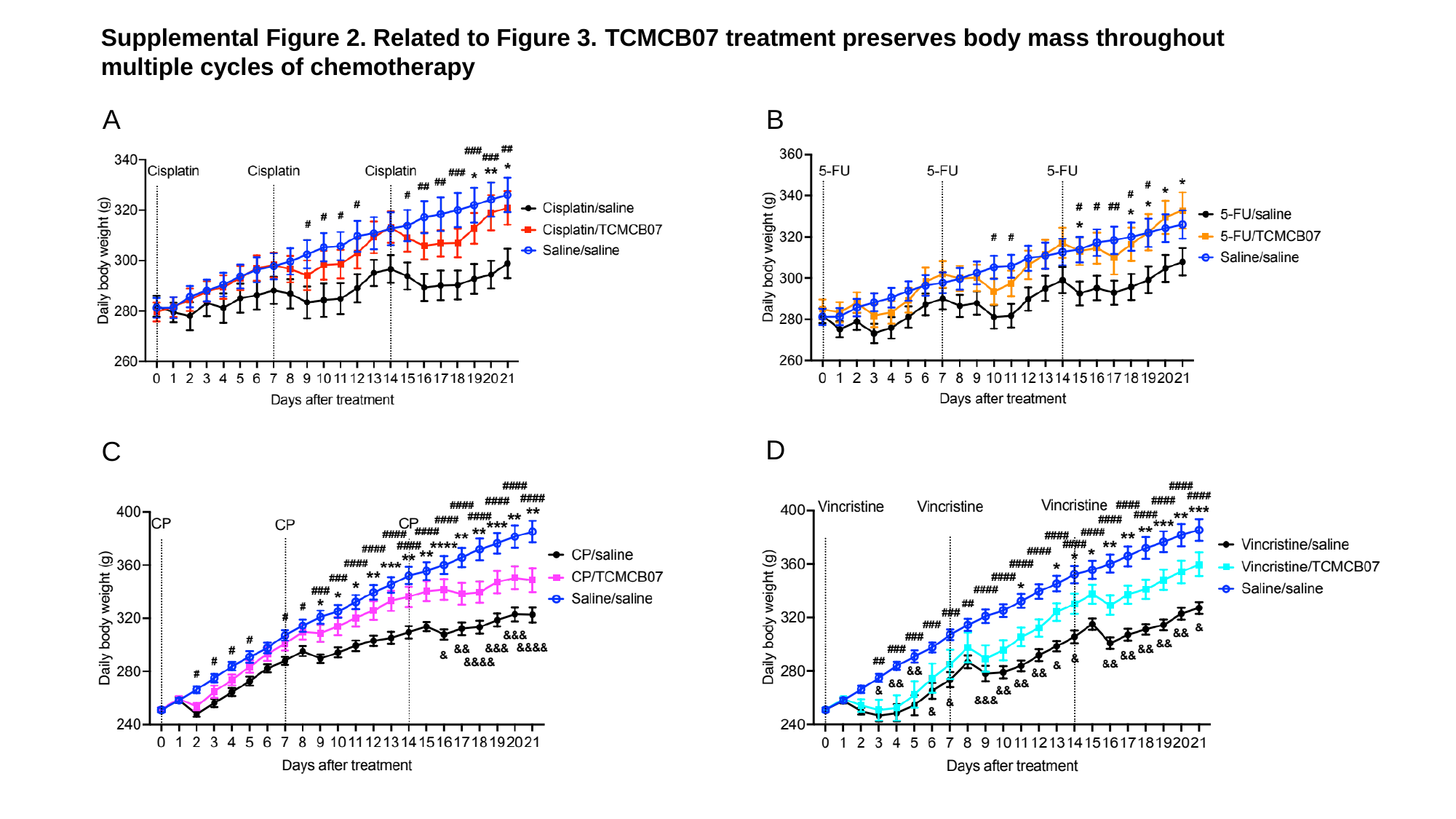

Supplemental Figure 2. Related to Figure 3. TCMCB07 treatment preserves body mass throughout multiple cycles of chemotherapy
A
B
D
C

### Slide 5
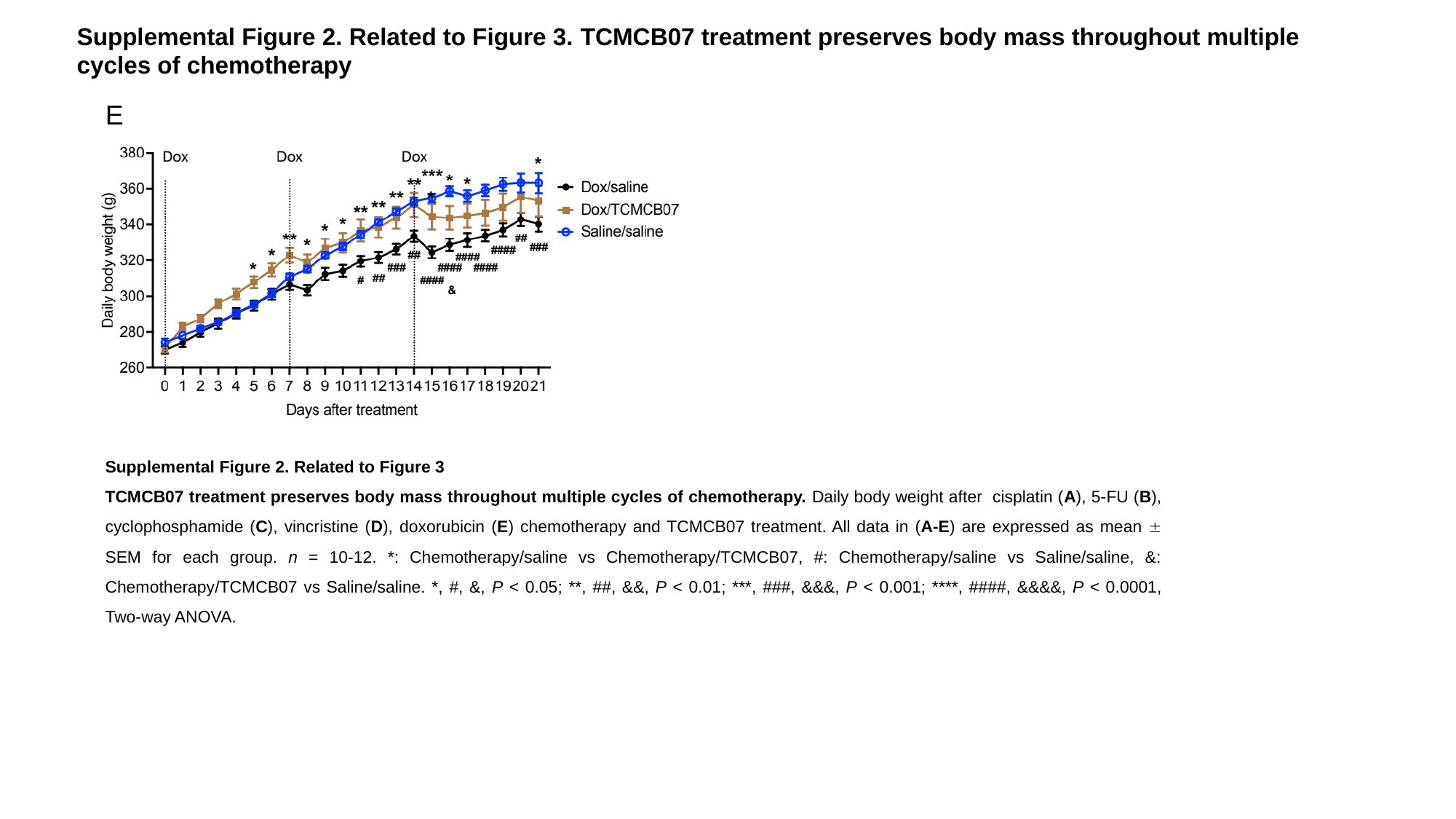

Supplemental Figure 2. Related to Figure 3. TCMCB07 treatment preserves body mass throughout multiple cycles of chemotherapy
E
Supplemental Figure 2. Related to Figure 3
TCMCB07 treatment preserves body mass throughout multiple cycles of chemotherapy. Daily body weight after cisplatin (A), 5-FU (B), cyclophosphamide (C), vincristine (D), doxorubicin (E) chemotherapy and TCMCB07 treatment. All data in (A-E) are expressed as mean ± SEM for each group. n = 10-12. *: Chemotherapy/saline vs Chemotherapy/TCMCB07, #: Chemotherapy/saline vs Saline/saline, &: Chemotherapy/TCMCB07 vs Saline/saline. *, #, &, P < 0.05; **, ##, &&, P < 0.01; ***, ###, &&&, P < 0.001; ****, ####, &&&&, P < 0.0001, Two-way ANOVA.

### Slide 6
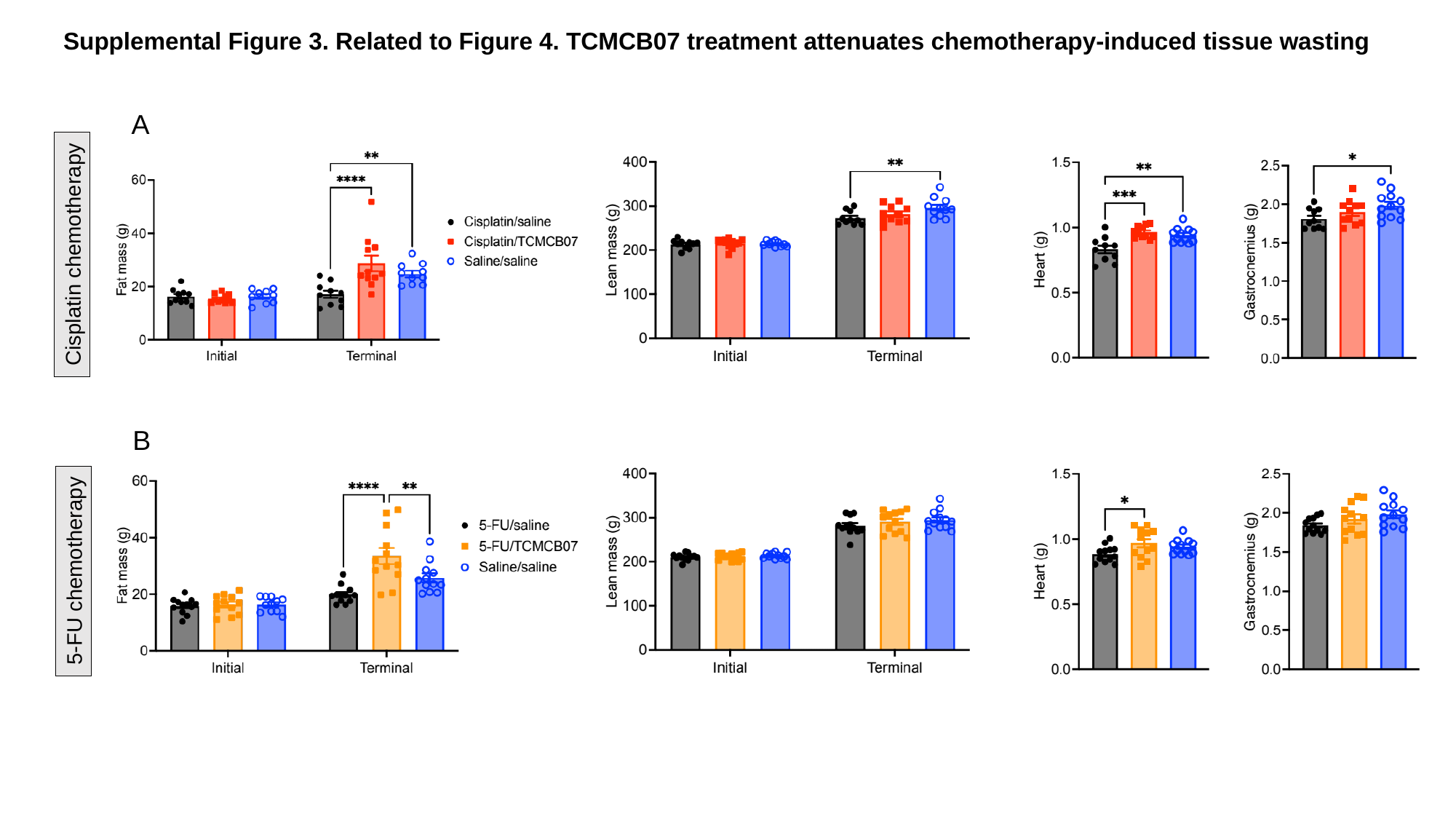

Supplemental Figure 3. Related to Figure 4. TCMCB07 treatment attenuates chemotherapy-induced tissue wasting
A
Cisplatin chemotherapy
B
5-FU chemotherapy

### Slide 7
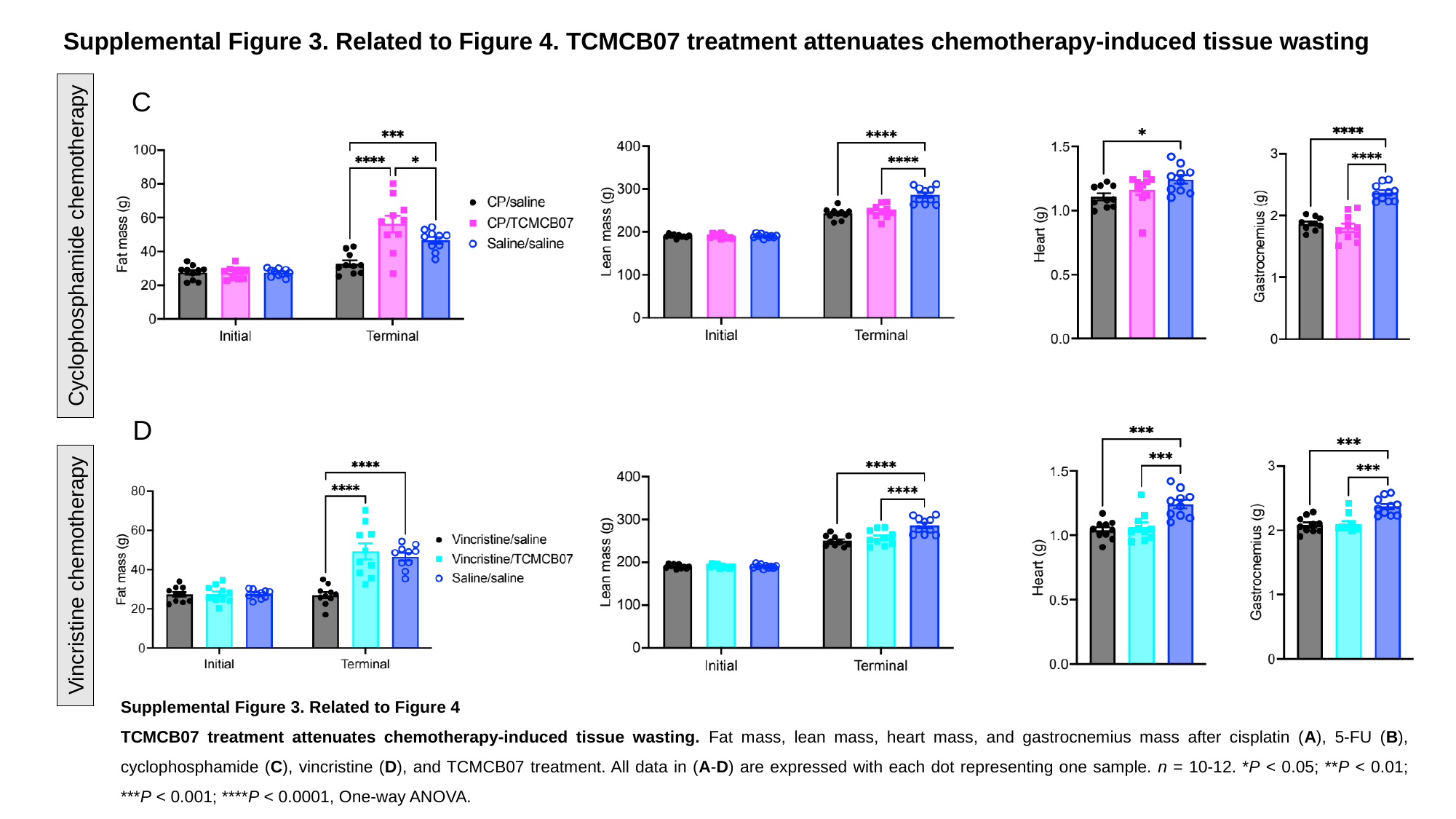

Supplemental Figure 3. Related to Figure 4. TCMCB07 treatment attenuates chemotherapy-induced tissue wasting
C
Cyclophosphamide chemotherapy
D
Vincristine chemotherapy
Supplemental Figure 3. Related to Figure 4
TCMCB07 treatment attenuates chemotherapy-induced tissue wasting. Fat mass, lean mass, heart mass, and gastrocnemius mass after cisplatin (A), 5-FU (B), cyclophosphamide (C), vincristine (D), and TCMCB07 treatment. All data in (A-D) are expressed with each dot representing one sample. n = 10-12. *P < 0.05; **P < 0.01; ***P < 0.001; ****P < 0.0001, One-way ANOVA.

### Slide 8
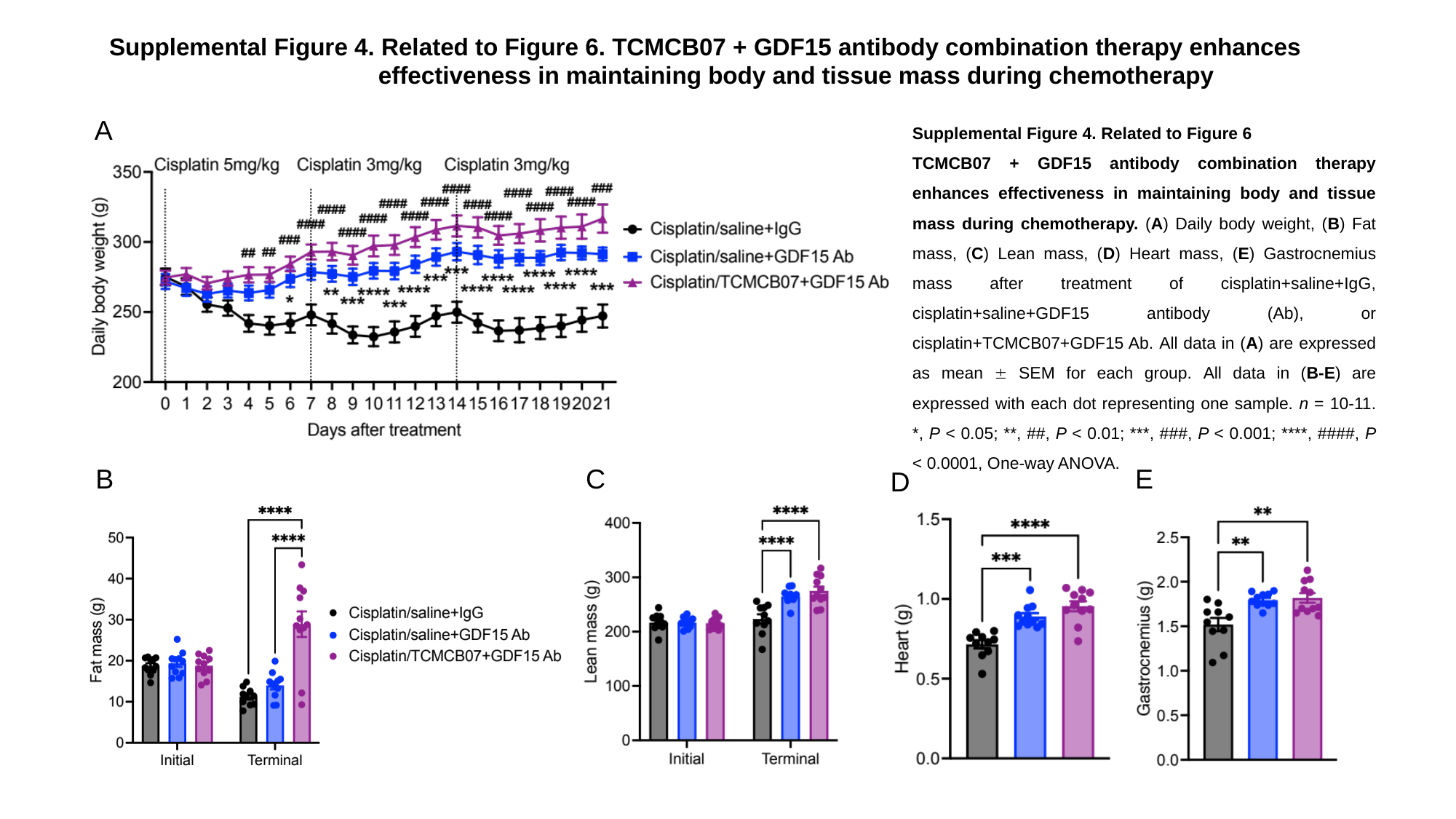

Supplemental Figure 4. Related to Figure 6. TCMCB07 + GDF15 antibody combination therapy enhances
 effectiveness in maintaining body and tissue mass during chemotherapy
A
Supplemental Figure 4. Related to Figure 6
TCMCB07 + GDF15 antibody combination therapy enhances effectiveness in maintaining body and tissue mass during chemotherapy. (A) Daily body weight, (B) Fat mass, (C) Lean mass, (D) Heart mass, (E) Gastrocnemius mass after treatment of cisplatin+saline+IgG, cisplatin+saline+GDF15 antibody (Ab), or cisplatin+TCMCB07+GDF15 Ab. All data in (A) are expressed as mean ± SEM for each group. All data in (B-E) are expressed with each dot representing one sample. n = 10-11. *, P < 0.05; **, ##, P < 0.01; ***, ###, P < 0.001; ****, ####, P < 0.0001, One-way ANOVA.
B
E
C
D

### Slide 9
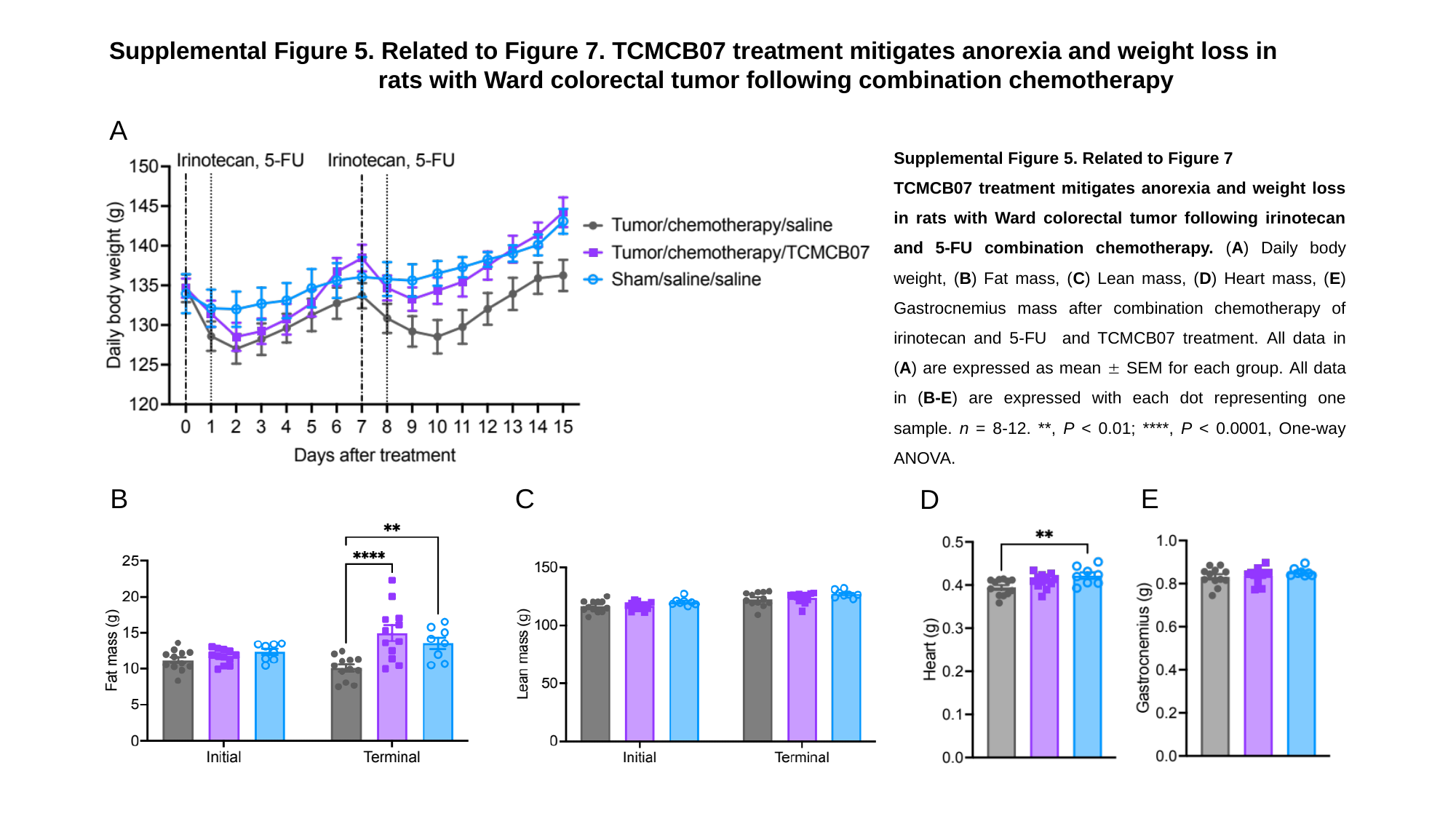

Supplemental Figure 5. Related to Figure 7. TCMCB07 treatment mitigates anorexia and weight loss in
 rats with Ward colorectal tumor following combination chemotherapy
A
Supplemental Figure 5. Related to Figure 7
TCMCB07 treatment mitigates anorexia and weight loss in rats with Ward colorectal tumor following irinotecan and 5-FU combination chemotherapy. (A) Daily body weight, (B) Fat mass, (C) Lean mass, (D) Heart mass, (E) Gastrocnemius mass after combination chemotherapy of irinotecan and 5-FU and TCMCB07 treatment. All data in (A) are expressed as mean ± SEM for each group. All data in (B-E) are expressed with each dot representing one sample. n = 8-12. **, P < 0.01; ****, P < 0.0001, One-way ANOVA.
B
C
E
D

### Slide 10
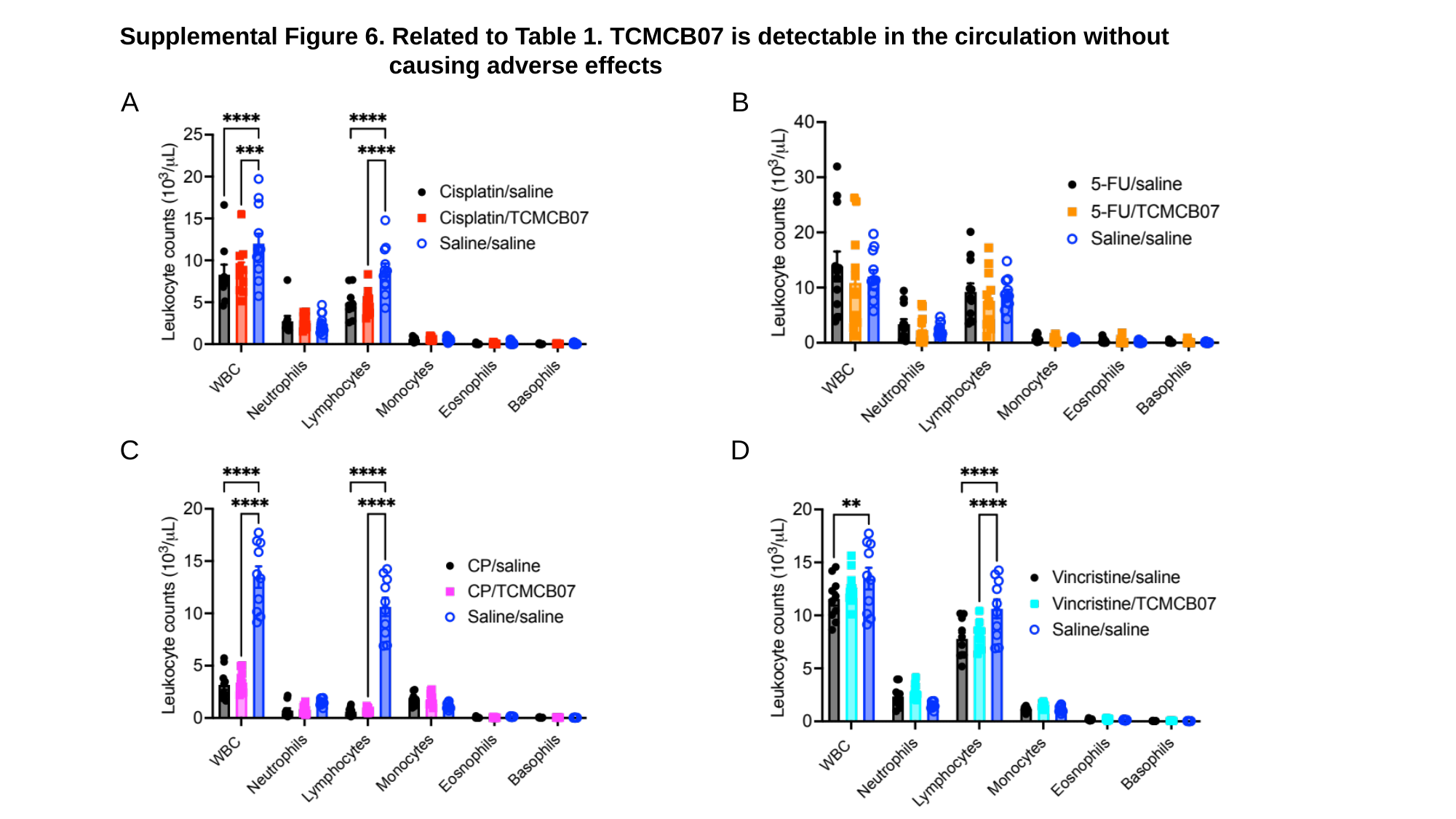

Supplemental Figure 6. Related to Table 1. TCMCB07 is detectable in the circulation without
 causing adverse effects
A
B
C
D

### Slide 11
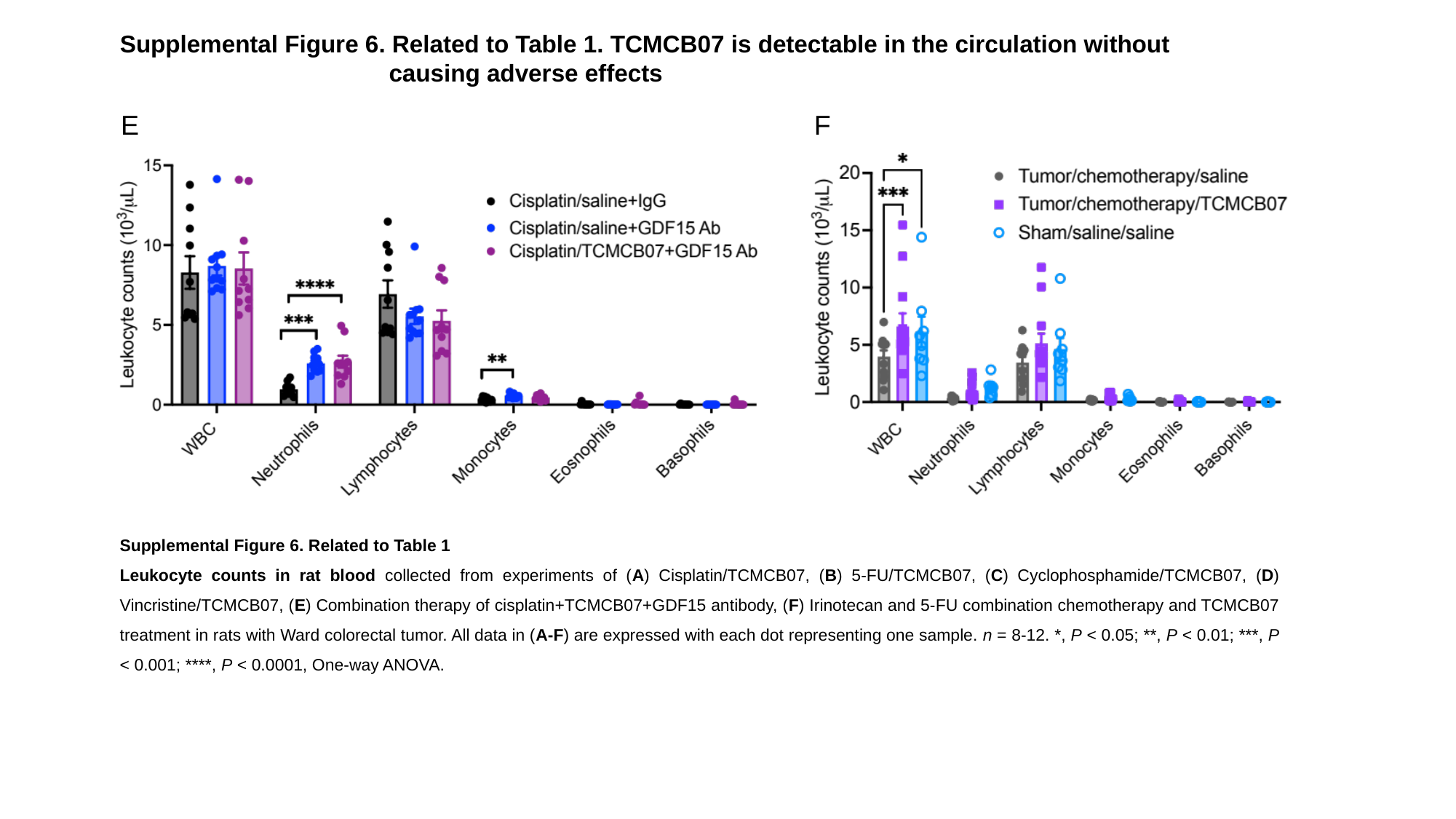

Supplemental Figure 6. Related to Table 1. TCMCB07 is detectable in the circulation without
 causing adverse effects
E
F
Supplemental Figure 6. Related to Table 1
Leukocyte counts in rat blood collected from experiments of (A) Cisplatin/TCMCB07, (B) 5-FU/TCMCB07, (C) Cyclophosphamide/TCMCB07, (D) Vincristine/TCMCB07, (E) Combination therapy of cisplatin+TCMCB07+GDF15 antibody, (F) Irinotecan and 5-FU combination chemotherapy and TCMCB07 treatment in rats with Ward colorectal tumor. All data in (A-F) are expressed with each dot representing one sample. n = 8-12. *, P < 0.05; **, P < 0.01; ***, P < 0.001; ****, P < 0.0001, One-way ANOVA.
